# On-demand, cell-free biomanufacturing of conjugate vaccines at the point-of-care

**DOI:** 10.1101/681841

**Authors:** Jessica C. Stark, Thapakorn Jaroentomeechai, Tyler D. Moeller, Rachel S. Dubner, Karen J. Hsu, Taylor C. Stevenson, Matthew P. DeLisa, Michael C. Jewett

## Abstract

Conjugate vaccines are among the most effective methods for preventing bacterial infections, representing a promising strategy to combat drug-resistant pathogens. However, existing manufacturing approaches limit access to conjugate vaccines due to centralized production and cold chain distribution requirements. To address these limitations, we developed a modular technology for *<u>i</u>n vitro* bioconjugate <u>va</u>ccine e<u>x</u>pression (iVAX) in portable, freeze-dried lysates from detoxified, nonpathogenic *Escherichia coli*. Upon rehydration, iVAX reactions synthesize clinically relevant doses of bioconjugate vaccines against diverse bacterial pathogens in one hour. We show that iVAX synthesized vaccines against the highly virulent pathogen *Franciscella tularensis* subsp. *tularensis* (type A) strain Schu S4 elicited pathogen-specific antibodies in mice at significantly higher levels compared to vaccines produced using engineered bacteria. The iVAX platform promises to accelerate development of new bioconjugate vaccines with increased access through refrigeration-independent distribution and point-of-care production.

## Introduction

Drug-resistant bacteria are predicted to threaten up to 10 million lives per year by 2050 (The Review on Antimicrobial Resistance, 2014), necessitating new strategies to develop and distribute antibiotics and vaccines. Conjugate vaccines, typically composed of a pathogen-specific capsular (CPS) or O-antigen polysaccharide (O-PS) linked to an immunostimulatory protein carrier, are among the safest and most effective methods for preventing life-threatening bacterial infections (Jin et al., 2017; Trotter et al., 2008; Weintraub, 2003). In particular, implementation of meningococcal and pneumococcal conjugate vaccines have significantly reduced the occurrence of bacterial meningitis and pneumonia worldwide (Novak et al., 2012; Poehling et al., 2006), in addition to reducing antibiotic resistance in targeted strains (Roush et al., 2008). However, despite their proven safety and efficacy, global childhood vaccination rates for conjugate vaccines remain as low as ~30%, with lack of access or low immunization coverage accounting for the vast majority of remaining disease burden (Wahl et al., 2018). In addition, the 2018 WHO prequalification of Typhbar-TCV^®^ to prevent typhoid fever represents the first conjugate vaccine approval in nearly a decade. In order to address emerging drug-resistant pathogens, new technologies to accelerate the development and global distribution of conjugate vaccines are needed.

Contributing to the slow pace of conjugate vaccine development and distribution is the fact that these molecules are particularly challenging and costly to manufacture. The conventional process to produce conjugate vaccines involves chemical conjugation of carrier proteins with polysaccharide antigens purified from large-scale cultures of pathogenic bacteria. Large-scale fermentation of pathogens results in high manufacturing costs due to associated biosafety hazards and process development challenges. In addition, chemical conjugation can alter the structure of the polysaccharide, resulting in loss of the protective epitope (Bhushan et al., 1998). To address these challenges, it was recently demonstrated that polysaccharide-protein “bioconjugates” can be made in *Escherichia coli* using protein-glycan coupling technology (PGCT) (Feldman et al., 2005). In this approach, engineered *E. coli* cells covalently attach heterologously expressed CPS or O-PS antigens to carrier proteins via an asparagine-linked glycosylation reaction catalyzed by the *Campylobacter jejuni* oligosaccharyltransferase enzyme PglB (*Cj*PglB) (Cuccui et al., 2013; Garcia-Quintanilla et al., 2014; Ihssen et al., 2010; Ma et al., 2014; Marshall et al., 2018; Wacker et al., 2014; Wetter et al., 2013). Despite this advance, both chemical conjugation and PGCT approaches rely on living bacterial cells, requiring centralized production facilities from which vaccines are distributed via a refrigerated supply chain. Refrigeration is critical to avoid conjugate vaccine spoilage due to precipitation and significant loss of the pathogen-specific polysaccharide upon both heating and freezing (Frasch, 2009; WHO, 2014). Only one conjugate vaccine, MenAfriVac™, is known to remain active outside of the cold chain for up to 4 days, which enabled increased vaccine coverage and an estimated 50% reduction in costs during vaccination in the meningitis belt of sub-Saharan Africa (Lydon et al., 2014). However, this required significant investment in the development and validation of a thermostable vaccine. Broadly, the need for cold chain refrigeration creates economic and logistical challenges that limit the reach of vaccination campaigns and present barriers to the eradication of disease, especially in the developing world (Ashok et al., 2017; Wahl et al., 2018).

Cell-free protein synthesis (CFPS) offers opportunities to both accelerate vaccine development and enable decentralized, cold chain-independent biomanufacturing by using cell lysates, rather than living cells, to synthesize proteins *in vitro* (Carlson et al., 2012). Importantly, CFPS platforms (i) enable point-of-care protein production, as relevant amounts of protein can be synthesized *in vitro* in just a few hours, (ii) can be freeze-dried for distribution at ambient temperature and reconstituted by just adding water (Adiga et al., 2018; Pardee et al., 2016), and (iii) circumvent biosafety concerns associated with the use of living cells outside of a controlled laboratory setting. CFPS has recently been used to enable on-demand and portable production of aglycosylated protein vaccines (Adiga et al., 2018; Pardee et al., 2016). Moreover, we recently described a cell-free glycoprotein synthesis technology that enables one-pot production of glycosylated proteins, including human glycoproteins and eukaryotic glycans (Jaroentomeechai et al., 2018). Despite these advances, cell-free systems and even decentralized manufacturing systems have historically been limited by their inability to synthesize glycosylated proteins at relevant titers and bearing diverse glycan structures, such as the polysaccharide antigens needed for bioconjugate vaccine production (Perez et al., 2016).

To address these limitations, here we describe the iVAX (*<u>i</u>n vitro* bioconjugate <u>va</u>ccine e<u>x</u>pression) platform that enables rapid development and cold chain-independent biosynthesis of conjugate vaccines in cell-free reactions (**Figure 1**). iVAX was designed to have the following features. First, iVAX is fast, with the ability to produce multiple individual doses of bioconjugates in one hour. Second, iVAX is robust, yielding equivalent amounts of bioconjugate over a range of operating temperatures. Third, iVAX is modular, offering the ability to rapidly interchange carrier proteins, including those used in licensed conjugate vaccines, as well as conjugated polysaccharide antigens. We leverage this modularity to create an array of vaccine candidates targeted against diverse bacterial pathogens, including the highly virulent *Franciscella tularensis* subsp. *tularensis* (type A) strain Schu S4, enterotoxigenic (ETEC) *E. coli* O78, and uropathogenic (UPEC) *E. coli* O7. Fourth, iVAX is shelf-stable, derived from freeze-dried cell-free reactions that operate in a just-add-water strategy. Fifth, iVAX is safe, leveraging lipid A remodeling that effectively avoids the high levels of endotoxin present in non-engineered *E. coli* manufacturing platforms. Our results demonstrate that anti-*F. tularensis* bioconjugates derived from freeze-dried, low-endotoxin iVAX reactions elicit pathogen-specific antibody responses in mice and outperform a bioconjugate produced using the established PGCT approach in living cells. Overall, the iVAX platform offers a new way to deliver the protective benefits of an important class of antibacterial vaccines to both the developed and developing world.

**Figure 1.**
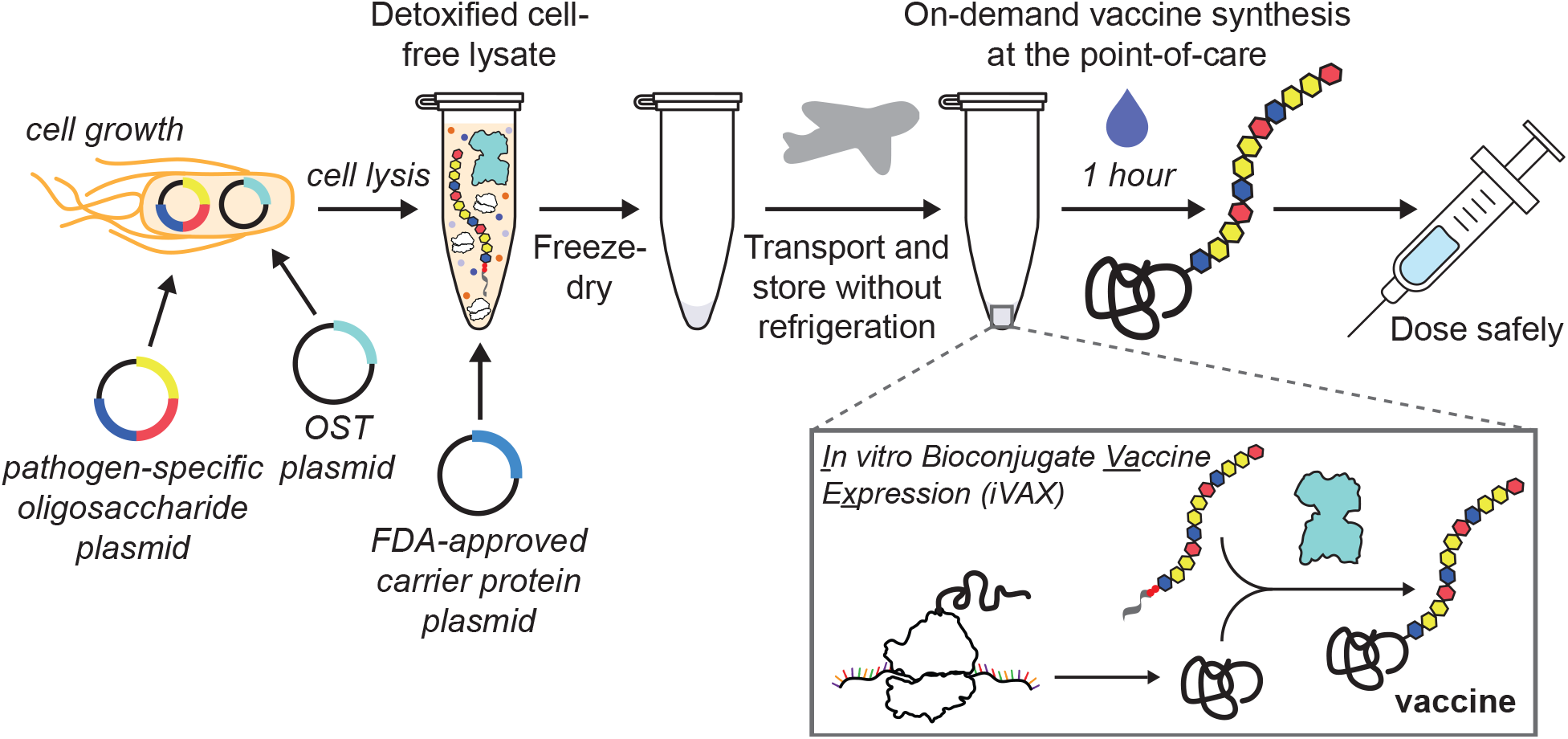
iVAX platform enables on-demand and portable production of antibacterial vaccines. The *<u>i</u>n vitro* bioconjugate <u>va</u>ccine e<u>x</u>pression (iVAX) platform provides a rapid means to develop and distribute vaccines against bacterial pathogens. Expression of pathogen-specific polysaccharides (e.g., CPS, O-PS) and a bacterial oligosaccharyltransferase enzyme in engineered nonpathogenic *E. coli* with detoxified lipid A yields low-endotoxin lysates containing all of the machinery required for synthesis of bioconjugate vaccines. Reactions catalyzed by iVAX lysates can be used to produce bioconjugates containing licensed carrier proteins and can be freeze-dried without loss of activity for refrigeration-free transportation and storage. Freeze-dried reactions can be activated at the point-of-care via simple rehydration and used to reproducibly synthesize immunologically active bioconjugates in ~1 h.

## Results

### *In vitro* synthesis of licensed vaccine carrier proteins

To demonstrate proof-of-principle for cell-free bioconjugate vaccine production, we first set out to express a set of carrier proteins that are currently used in FDA-approved conjugate vaccines. Producing these carrier proteins in soluble conformations *in vitro* represented an important benchmark because their expression in living *E. coli* has proven challenging, often requiring purification and refolding of insoluble product from inclusion bodies (Haghi et al., 2011; Stefan et al., 2011), fusion of expression partners such as maltose-binding protein (MBP) to increase soluble expression (Figueiredo et al., 1995; Stefan et al., 2011), or expression of truncated protein variants in favor of the full-length proteins (Figueiredo et al., 1995). In contrast, cell-free protein synthesis approaches have recently shown promise for difficult-to-express proteins (Perez et al., 2016). The carrier proteins that we focused on here included nonacylated *H. influenzae* protein D (PD), the *N. meningitidis* porin protein (PorA), and genetically detoxified variants of the *Corynebacterium diphtheriae* toxin (CRM197) and the *Clostridium tetani* toxin (TT). We also tested expression of the fragment C (TTc) and light chain (TTlight) domains of TT as well as *E. coli* MBP. While MBP is not a licensed carrier, it has demonstrated immunostimulatory properties (Fernandez et al., 2007) and when linked to O-PS was found to elicit polysaccharide-specific humoral and cellular immune responses in mice (Ma et al., 2014). Similarly, the TT domains, TTlight and TTc, have not been used in licensed vaccines, but are immunostimulatory and individually sufficient for protection against *C. tetani* challenge in mice (Figueiredo et al., 1995). To enable glycosylation, all carriers were modified at their C-termini with 4 tandem repeats of an optimal bacterial glycosylation motif, DQNAT (Chen et al., 2007). A C-terminal 6xHis tag was also included to enable purification and detection via Western blot analysis. A variant of superfolder green fluorescent protein that contained an internal DQNAT glycosylation site (sfGFP^217-DQNAT^) (Jaroentomeechai et al., 2018) was used as a model protein to facilitate system development.

All eight carriers were synthesized *in vitro* with soluble yields of ~50-650 μg mL^−1^ as determined by ^14^C-leucine incorporation (**Figure 2a**). In particular, the MBP^4xDQNAT^ and PD^4xDQNAT^ variants were nearly 100% soluble, with yields of 500 μg mL^−1^ and 200 μg mL^−1^, respectively, and expressed as exclusively full-length products according to Western blot analysis (**Figure 2b**). Notably, similar soluble yields were observed for all carriers at 25°C, 30°C, and 37°C, with the exception of CRM197^4xDQNAT^ (**Figure S1a**), which is known to be heat sensitive (WHO, 2014). These results suggest that our method of cell-free carrier biosynthesis is robust over a 13°C range in temperature and could be used in settings where precise temperature control is not feasible.

**Figure 2.**
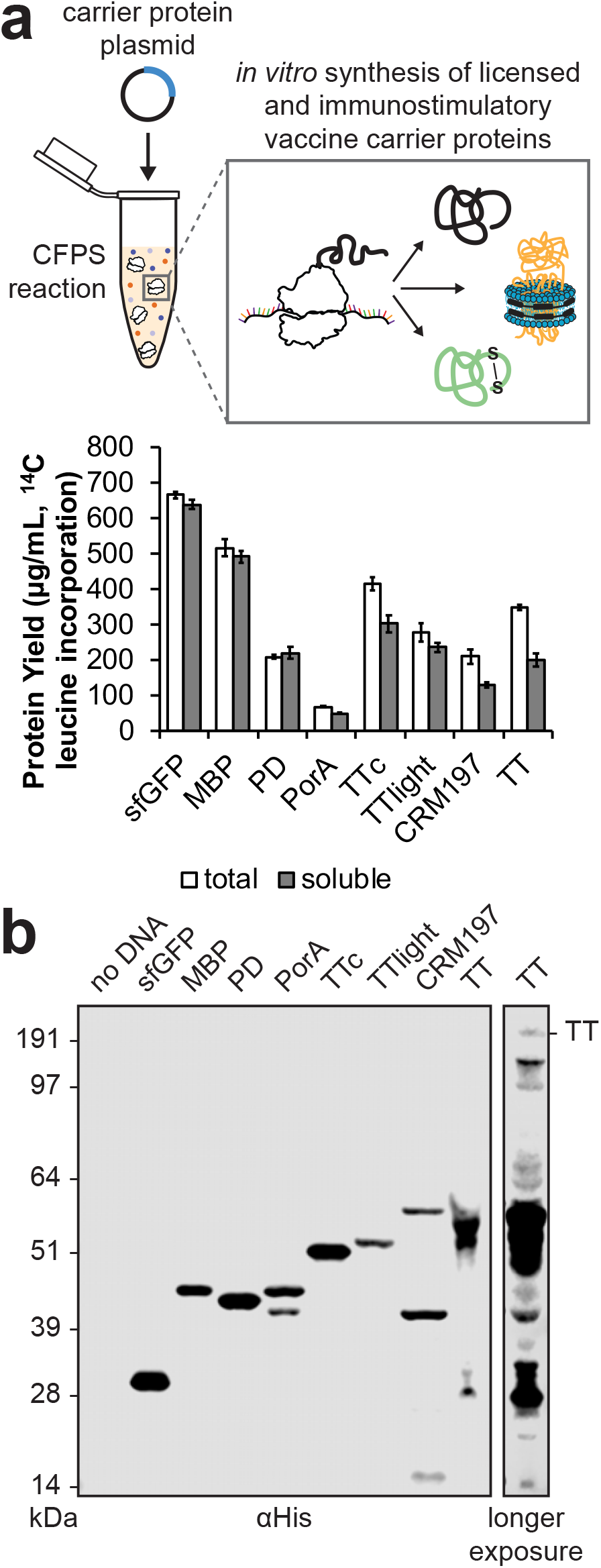
*In vitro* synthesis of licensed conjugate vaccine carrier proteins. (**a**) All four carrier proteins used in FDA-approved conjugate vaccines were synthesized solubly *in vitro*, as measured via ^14^C-leucine incorporation. These include *H. influenzae* protein D (PD), the *N. meningitidis* porin protein (PorA), and genetically detoxified variants of the *C. diphtheriae* toxin (CRM197) and the *C. tetani* toxin (TT). Additional immunostimulatory carriers were also synthesized solubly, including *E. coli* maltose binding protein (MBP) and the fragment C (TTc) and light chain (TTlight) domains of TT. Values represent means and error bars represent standard deviations of biological replicates (*n* = 3). (**b**) Full length product was observed for all proteins tested via Western blot. Different exposures are indicated with solid lines. Molecular weight ladder is shown at left. *See also Figure S1.*

The open reaction environment of our cell-free reactions enabled facile manipulation of the chemical and reaction environment to improve production of more complex carriers. For example, in the case of the membrane protein PorA^4xDQNAT^, lipid nanodiscs were added to increase soluble expression (**Figure S1b**). Nanodiscs provide a cellular membrane mimic to co-translationally stabilize hydrophobic regions of membrane proteins (Bayburt and Sligar, 2010). For expression of TT, which contains an intermolecular disulfide bond, expression was carried out for 2 hours in oxidizing conditions (Knapp et al., 2007), which improved assembly of the heavy and light chains into full-length product and minimized protease degradation of full-length TT (**Figure S1c**). *In vitro* synthesized CRM197^4xDQNAT^ and TT^4xDQNAT^ were comparable in size to commercially available purified diphtheria toxin (DT) and TT protein standards and were reactive with α-DT and α-TT antibodies, respectively (**Figure S1d, e**), indicating that both were produced in immunologically relevant conformations. This is notable as CRM197 and TT are FDA-approved vaccine antigens for diphtheria and tetanus, respectively, when they are administered without conjugated polysaccharides. Together, our results highlight the ability of CFPS to express licensed conjugate vaccine carrier proteins in soluble conformations over a range of temperatures.

### On-demand synthesis of bioconjugate vaccines

We next sought to synthesize polysaccharide-conjugated versions of these carrier proteins by merging their *in vitro* expression with one-pot, cell-free glycosylation. As a model vaccine target, we focused on the highly virulent *Francisella tularensis* subsp. *tularensis* (type A) strain Schu S4, a gram-negative, facultative coccobacillus and the causative agent of tularemia. This bacterium is categorized as a class A bioterrorism agent due to its high fatality rate, low dose of infection, and ability to be aerosolized (Oyston et al., 2004). Although there are currently no licensed vaccines against *F. tularensis*, several studies have independently confirmed the important role of antibodies directed against *F. tularensis* LPS, specifically the O-PS repeat unit, in providing protection against the Schu S4 strain (Fulop et al., 2001; Lu et al., 2012). More recently, a bioconjugate vaccine comprising the *F. tularensis* Schu S4 O-PS (*Ft*O-PS) conjugated to the *Pseudomonas aeruginosa* exotoxin A (EPA^DNNNS-DQNRT^) carrier protein produced using PGCT (Cuccui et al., 2013; Marshall et al., 2018) was shown to be protective against challenge with the Schu S4 strain in a rat inhalation model of tularemia (Marshall et al., 2018). In light of these earlier findings, we investigated the ability of the iVAX platform to produce anti-*F. tularensis* bioconjugate vaccine candidates on-demand by conjugating the *Ft*O-PS structure to diverse carrier proteins *in vitro*.

The *Ft*O-PS is composed of the 826-Da repeating tetrasaccharide unit Qui4NFm-(GalNAcAN)_2_-QuiNAc (Qui4NFm: 4,6-dideoxy-4-formamido-D-glucose; GalNAcAN: 2-acetamido-2-deoxy-D-galacturonamide; QuiNAc: 2-acetamido-2,6-dideoxy-D-glucose) (Prior et al., 2003). To glycosylate proteins with *Ft*O-PS, we produced an iVAX lysate from glycoengineered *E. coli* cells expressing the *Ft*O-PS biosynthetic pathway and the oligosaccharyltransferase enzyme *Cj*PglB (**Figure 3a**). This lysate, which contained lipid-linked *Ft*O-PS and active *Cj*PglB, was used to catalyze iVAX reactions primed with plasmid DNA encoding sfGFP^217-DQNAT^. Control reactions in which attachment of the *Ft*O-PS was not expected were performed with lysates from cells that lacked either the *Ft*O-PS pathway or the *Cj*PglB enzyme. We also tested reactions that lacked plasmid encoding the target protein sfGFP^217-DQNAT^ or were primed with plasmid encoding sfGFP^217-AQNAT^, which contained a mutated glycosylation site (AQNAT) that is not modified by *Cj*PglB (Kowarik et al., 2006). In reactions containing the iVAX lysate and primed with plasmid encoding sfGFP^217-DQNAT^, immunoblotting with anti-His antibody or a commercial monoclonal antibody specific to *Ft*O-PS revealed a ladder-like banding pattern (**Figure 3b**). This ladder is characteristic of *Ft*O-PS attachment, resulting from O-PS chain length variability through the action of the Wzy polymerase (Cuccui et al., 2013; Feldman et al., 2005; Prior et al., 2003). Glycosylation of sfGFP^217-DQNAT^ was observed only in reactions containing a complete glycosylation pathway and the preferred DQNAT glycosylation sequence (**Figure 3b**). This glycosylation profile was reproducible across biological replicates from the same lot of lysate (**Figure 3c**, **left**) and using different lots of lysate (**Figure 3c**, **right**). *In vitro* protein synthesis and glycosylation was observed after 1 hour, with the amount of conjugated polysaccharide reaching a maximum between 0.75 and 1.25 hours (**Figure S2**). Similar glycosylation reaction kinetics were observed at 37°C, 30°C, 25°C, and room temperature (~21°C), indicating that iVAX reactions are robust over a range of temperatures (**Figure S2**).

**Figure 3.**
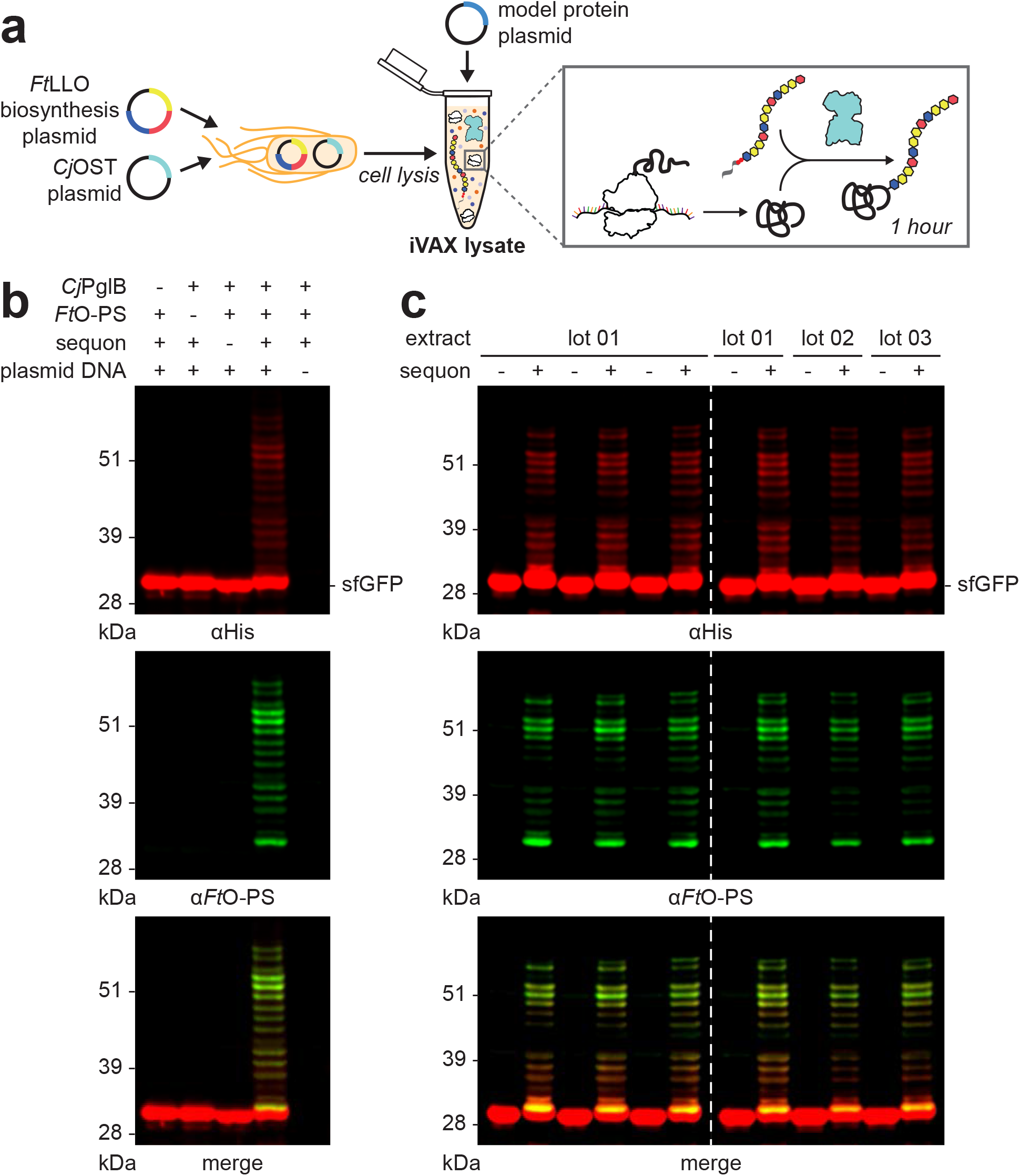
Reproducible glycosylation of proteins with *Ft*O-PS in iVAX lysates. (**a**) iVAX lysates were prepared from cells expressing *Cj*PglB and a biosynthetic pathway encoding *Ft*O-PS. (**b**) Glycosylation of sfGFP^217-DQNAT^ with *Ft*O-PS was only observed when *Cj*PglB, *Ft*O-PS, and the preferred sequon were present in the reaction (lane 3). When plasmid DNA was omitted, sfGFP^217-DQNAT^ synthesis was not observed. (**c**) Biological replicates of iVAX reactions producing sfGFP^217-DQNAT^ using the same lot (**left**) or different lots (**right**) of iVAX lysates demonstrated reproducibility of reactions and lysate preparation. Top panels show signal from probing with anti-hexa-histidine antibody (αHis) to detect the carrier protein, middle panels show signal from probing with commercial anti-*Ft*O-PS antibody (α*Ft*O-PS), and bottom panels show αHis and α*Ft*O-PS signals merged. Unless replicates are explicitly shown, images are representative of at least three biological replicates. Dashed lines indicate samples are from the same blot with the same exposure. Molecular weight ladders are shown at the left of each image. *See also Figure S2.*

Next, we investigated whether FDA-approved carriers could be similarly conjugated with *Ft*O-PS in iVAX reactions. Following addition of plasmid DNA encoding MBP^4xDQNAT^, PD^4xDQNAT^, PorA^4xDQNAT^, TTc^4xDQNAT^, TTlight^4xDQNAT^, and CRM197^4xDQNAT^, glycosylation of each with *Ft*O-PS was observed for iVAX reactions enriched with lipid-linked *Ft*O-PS and *Cj*PglB but not control reactions lacking *Cj*PglB (**Figure 4**). We observed conjugation of high molecular weight *Ft*O-PS species (on the order of ~10-20 kDa) to all protein carriers tested, which is important as longer glycan chain length has been shown to increase the efficacy of conjugate vaccines against *F. tularensis* (Stefanetti et al., 2019). Notably, our attempts to synthesize the same panel of bioconjugates using the established PGCT approach in living *E. coli* yielded less promising results. Specifically, we observed low levels of *Ft*O-PS glycosylation and lower molecular weight conjugated *Ft*O-PS species for all PGCT-derived bioconjugates compared to their iVAX-derived counterparts (**Figure S3**). The same trend was observed for PGCT- versus iVAX-derived bioconjugates involving the most common PGCT carrier protein, EPA^DNNNS-DQNRT^ (Cuccui et al., 2013; Ihssen et al., 2010; Marshall et al., 2018; Wacker et al., 2014; Wetter et al., 2013) (**Figure 4**; **Figure S3**). In addition, only limited expression of the PorA membrane protein was achieved *in vivo* (**Figure S3**). Collectively, these data indicate that iVAX could provide advantages over PGCT for production of bioconjugate vaccine candidates with high molecular weight O-PS antigens conjugated efficiently to diverse and potentially membrane-bound carrier proteins.

**Figure 4.**
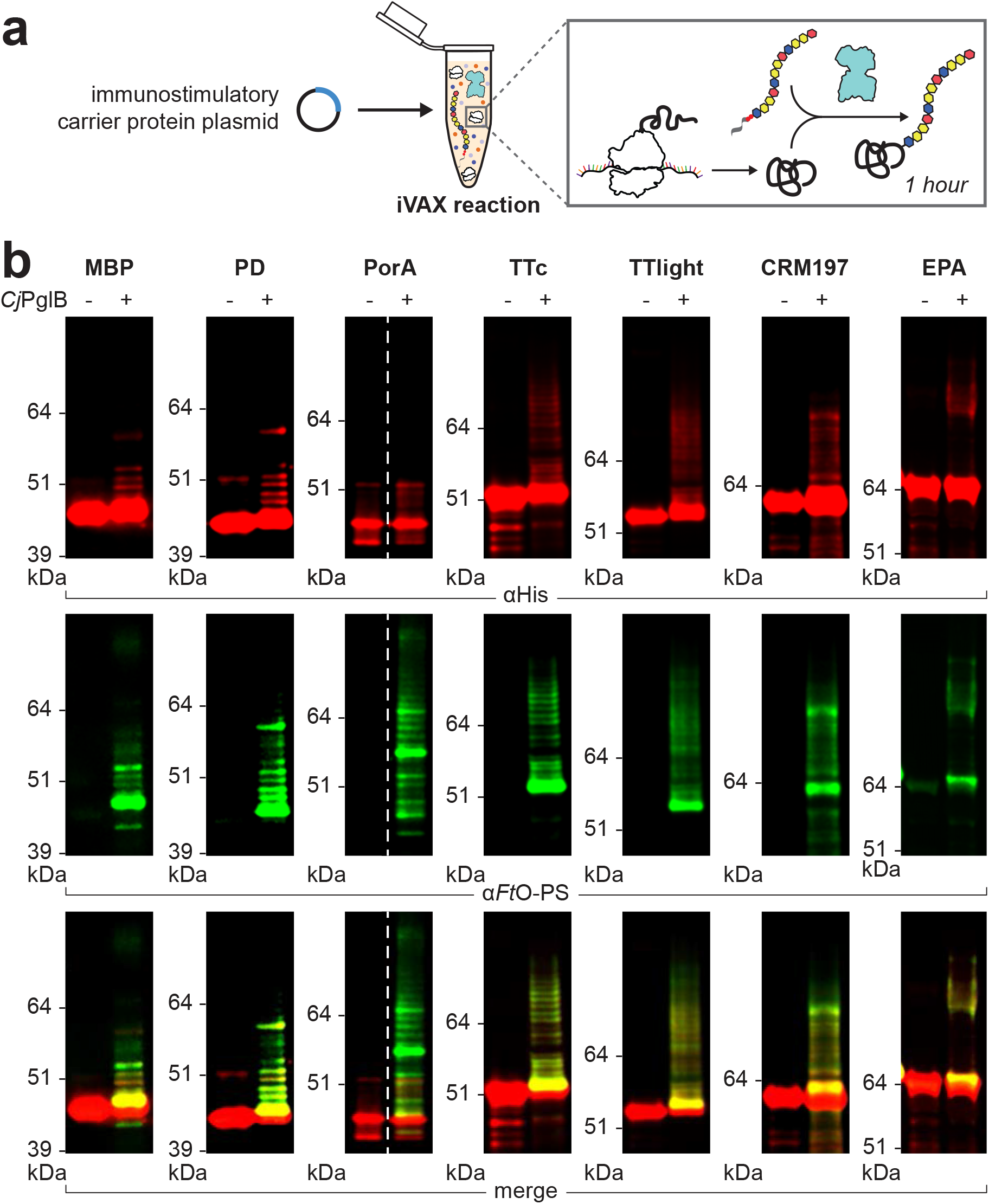
On-demand production of bioconjugates against *F. tularensis* using iVAX. (**a**) iVAX reactions were prepared from lysates containing *Cj*PglB and *Ft*O-PS and primed with plasmid encoding immunostimulatory carriers, including those used in licensed vaccines. (**b**) We observed on-demand synthesis of anti-*F. tularensis* bioconjugate vaccines for all carrier proteins tested. Bioconjugates were purified using Ni-NTA agarose from 1 mL iVAX reactions lasting ~1 h. Top panels show signal from probing with anti-hexa-histidine antibody (αHis) to detect the carrier protein, middle panels show signal from probing with commercial anti-*Ft*O-PS antibody (α*Ft*O-PS), and bottom panels show αHis and α*Ft*O-PS signals merged. Images are representative of at least three biological replicates. Dashed lines indicate samples are from the same blot with the same exposure. Molecular weight ladders are shown at the left of each image. *See also Figures S3 and S4.*

We next asked whether the yields of bioconjugates produced using iVAX were sufficient to enable production of relevant vaccine doses. Recent clinical data show 1-10 μg doses of bioconjugate vaccine candidates are well-tolerated and effective in stimulating the production of antibacterial IgGs (Hatz et al., 2015; Huttner et al., 2017; Riddle et al., 2016). To assess expression titers and for future experiments, we focused on MBP^4xDQNAT^ and PD^4xDQNAT^ because these carriers expressed *in vitro* with high soluble titers and without truncation products (**Figure 2**). We found that reactions lasting ~1 hour produced ~20 μg mL^−1^ of glycosylated MBP^4xDQNAT^ and PD^4xDQNAT^ as determined by ^14^C-leucine incorporation and densitometry analysis (**Figure S4a**). It should be noted that vaccines are currently distributed in vials containing 1-20 doses of vaccine to minimize wastage (Humphreys, 2011). Our yields suggest that multiple doses per mL can be synthesized in 1 hour using the iVAX platform.

To demonstrate the modularity of the iVAX approach for bioconjugate production, we sought to produce bioconjugates bearing O-PS antigens from additional pathogens including ETEC *E. coli* strain O78 and UPEC *E. coli* strain O7. *E. coli* O78 is a major cause of diarrheal disease in developing countries, especially among children, and a leading cause of traveler’s diarrhea (Qadri et al., 2005), while the O7 strain is a common cause of urinary tract infections (Johnson, 1991). Like the *Ft*O-PS, the biosynthetic pathways for *Ec*O78-PS and *Ec*O7-PS have been described previously and confirmed to produce O-PS antigens with the repeating units GlcNAc_2_Man_2_ (Jansson et al., 1987) and Qui4NAcMan(Rha)GalGlcNAc (L’vov et al., 1984) (GlcNAc: *N*-acetylglucosamine; Man: mannose; Qui4NAc: 4-acetamido-4,6-dideoxy-D-glucopyranose; Rha: rhamnose; Gal: galactose), respectively. Using iVAX lysates from cells expressing *Cj*PglB and either the *Ec*O78-PS and *Ec*O7-PS pathways in reactions that were primed with PD^4xDQNAT^ or sfGFP^217-DQNAT^ plasmids, we observed carrier glycosylation only when both lipid-linked O-PS and *Cj*PglB were present in the reactions (**Figure S4b, c**). Our results demonstrate modular production of bioconjugates against multiple bacterial pathogens, enabled by compatibility of multiple heterologous O-PS pathways with *in vitro* carrier protein synthesis and glycosylation.

### Endotoxin editing and freeze-drying yield iVAX reactions that are safe and portable

A key challenge inherent in using any *E. coli*-based system for biopharmaceutical production is the presence of lipid A, or endotoxin, which is known to contaminate protein products and can cause lethal septic shock at high levels (Russell, 2006). As a result, the amount of endotoxin in formulated biopharmaceuticals is regulated by the United States Pharmacopeia (USP), US Food and Drug Administration (FDA), and the European Medicines Agency (EMEA) (Brito and Singh, 2011). Because our iVAX reactions rely on lipid-associated components, such as *Cj*PglB and *Ft*O-PS, standard detoxification approaches involving the removal of lipid A (Petsch and Anspach, 2000) could compromise the activity or concentration of our glycosylation components in addition to increasing cost and processing complexities.

To address this issue, we adapted a previously reported strategy to detoxify the lipid A molecule through strain engineering (Chen et al., 2016; Needham et al., 2013). In particular, the deletion of the acyltransferase gene *lpxM* and the overexpression of the *F. tularensis* phosphatase LpxE in *E. coli* has been shown to result in the production of nearly homogenous pentaacylated, monophosphorylated lipid A with significantly reduced toxicity but retained activity as an adjuvant (Chen et al., 2016). This pentaacylated, monophosphorylated lipid A was structurally identical to the primary component of monophosphoryl lipid A (MPL) from *Salmonella minnesota* R595, an approved adjuvant composed of a mixture of monophosphorylated lipids (Casella and Mitchell, 2008). To generate detoxified lipid A structures in the context of iVAX, we produced lysates from a Δ*lpxM* derivative of CLM24 that co-expressed *Ft*LpxE and the *Ft*O-PS glycosylation pathway (**Figure 5a**). Lysates derived from this strain exhibited significantly decreased levels of toxicity compared to wild type CLM24 lysates expressing *Cj*PglB and *Ft*O-PS (**Figure 5b**) as measured by human TLR4 activation in HEK-Blue hTLR4 reporter cells (Needham et al., 2013). Importantly, the structural remodeling of lipid A did not affect the activity of the membrane-bound *Cj*PglB and *Ft*O-PS components in iVAX reactions (**Figure S5a**). By engineering the chassis strain for lysate production, we produced iVAX lysates with endotoxin levels <1,000 EU mL^−1^, within the range of reported values for commercial protein-based vaccine products (0.288-180,000 EU mL^−1^) (Brito and Singh, 2011).

**Figure 5.**
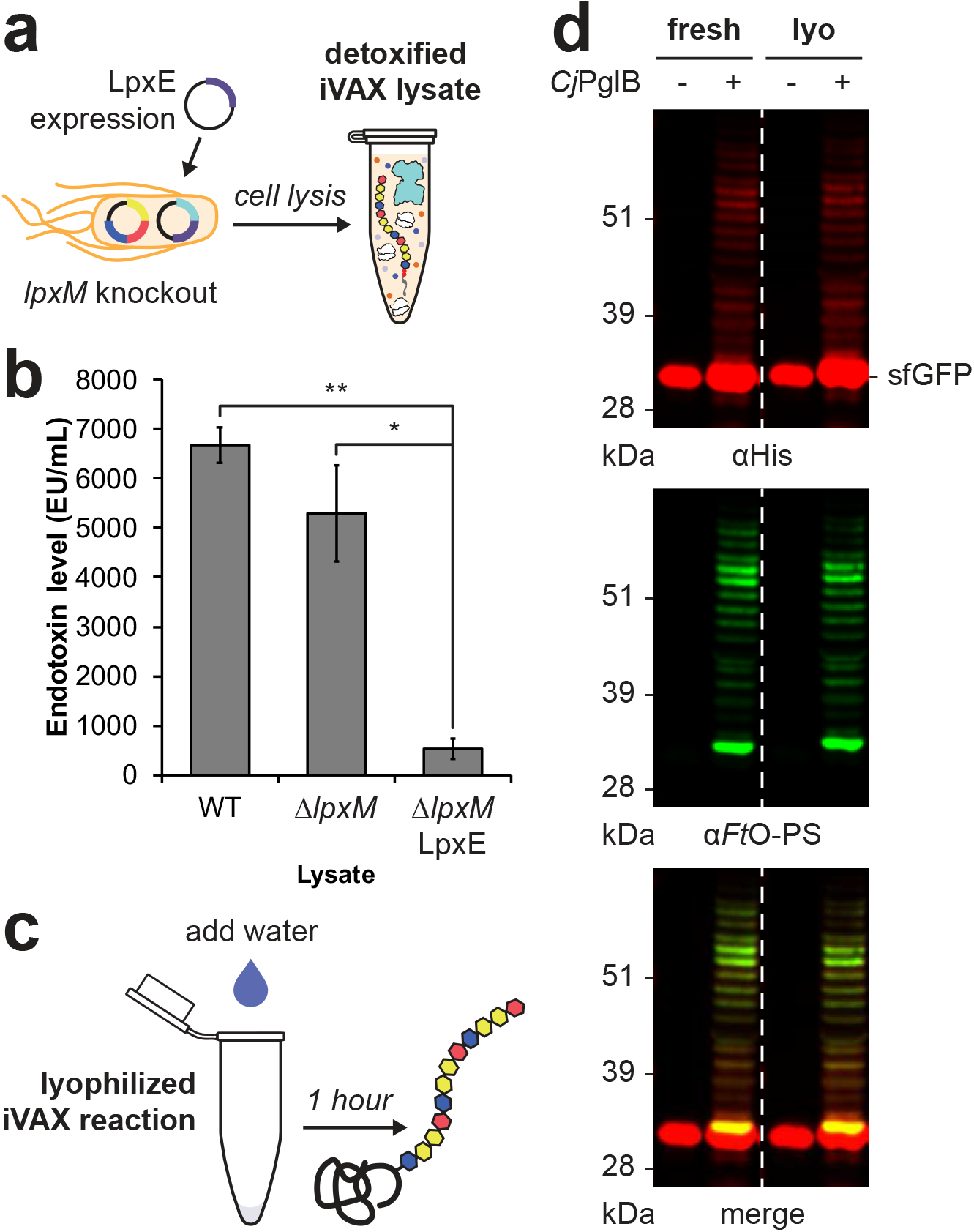
Detoxified, lyophilized iVAX reactions produce bioconjugates. (**a**) iVAX lysates were detoxified via deletion of *lpxM* and expression of *F. tularensis* LpxE in the source strain for lysate production. (**b**) The resulting lysates exhibited significantly reduced endotoxin activity. \**p* = 0.019 and \*\**p* = 0.003, as determined by two-tailed *t*-test. (**c**) iVAX reactions producing sfGFP^217-DQNAT^ were run immediately or following lyophilization and rehydration. (**d**) Glycosylation activity was preserved following lyophilization, demonstrating the potential of iVAX reactions for portable biosynthesis of bioconjugate vaccines. Top panel shows signal from probing with anti-hexa-histidine antibody (αHis) to detect the carrier protein, middle panel shows signal from probing with commercial anti-*Ft*O-PS antibody (α*Ft*O-PS), and bottom panel shows αHis and α*Ft*O-PS signals merged. Images are representative of at least three biological replicates. Molecular weight ladder is shown at the left of each image. *See also Figure S5.*

A major limitation of traditional conjugate vaccines is that they must be refrigerated (WHO, 2014), making it difficult to distribute these vaccines to remote or resource-limited settings. The ability to freeze-dry iVAX reactions for ambient temperature storage and distribution could alleviate the logistical challenges associated with refrigerated supply chains that are required for existing vaccines. To investigate this possibility, detoxified iVAX lysates were used to produce *Ft*O-PS bioconjugates in two different ways: either by running the reaction immediately after priming with plasmid encoding the sfGFP^217-DQNAT^ target protein or by running after the same reaction mixture was lyophilized and rehydrated (**Figure 5c**). In both cases, conjugation of *Ft*O-PS to sfGFP^217-DQNAT^ was observed when *Cj*PglB was present, with modification levels that were nearly identical (**Figure 5d**). We also showed that detoxified, freeze-dried iVAX reactions can be scaled to 5 mL for production of *Ft*O-PS-conjugated MBP^4xDQNAT^ and PD^4xDQNAT^ in a manner that was reproducible from lot to lot and indistinguishable from production without freeze-drying (**Figure S5b, c**). The ability to lyophilize iVAX reactions and manufacture bioconjugates without specialized equipment highlights the potential for portable, on-demand vaccine production.

### *In vitro* synthesized bioconjugates elicit pathogen-specific antibodies in mice

To validate the efficacy of bioconjugates produced using the iVAX platform, we next evaluated the ability of iVAX-derived bioconjugates to elicit anti-*Ft*LPS antibodies in mice (**Figure 6a**). Importantly, we found that BALB/c mice receiving iVAX-derived *Ft*O-PS-conjugated MBP^4xDQNAT^ or PD^4xDQNAT^ produced high titers of *Ft*LPS-specific IgG antibodies, which were significantly elevated compared to the titers measured in the sera of control mice receiving PBS or aglycosylated versions of each carrier protein (**Figure 6b**, **Figure S6**). Interestingly, the IgG titers measured in sera from mice receiving glycosylated MBP^4xDQNAT^ derived from PGCT were similar to the titers observed in the control groups (**Figure 6b**, **Figure S6**), in line with the markedly weaker glycosylation of this candidate relative to its iVAX-derived counterpart (**Figure S3**). Notably, both MBP^4xDQNAT^ and PD^4xDQNAT^ bioconjugates produced using iVAX elicited similar levels of IgG production and neither resulted in any observable adverse events in mice, confirming the modularity and safety of the technology for production of bioconjugate vaccine candidates.

**Figure 6.**
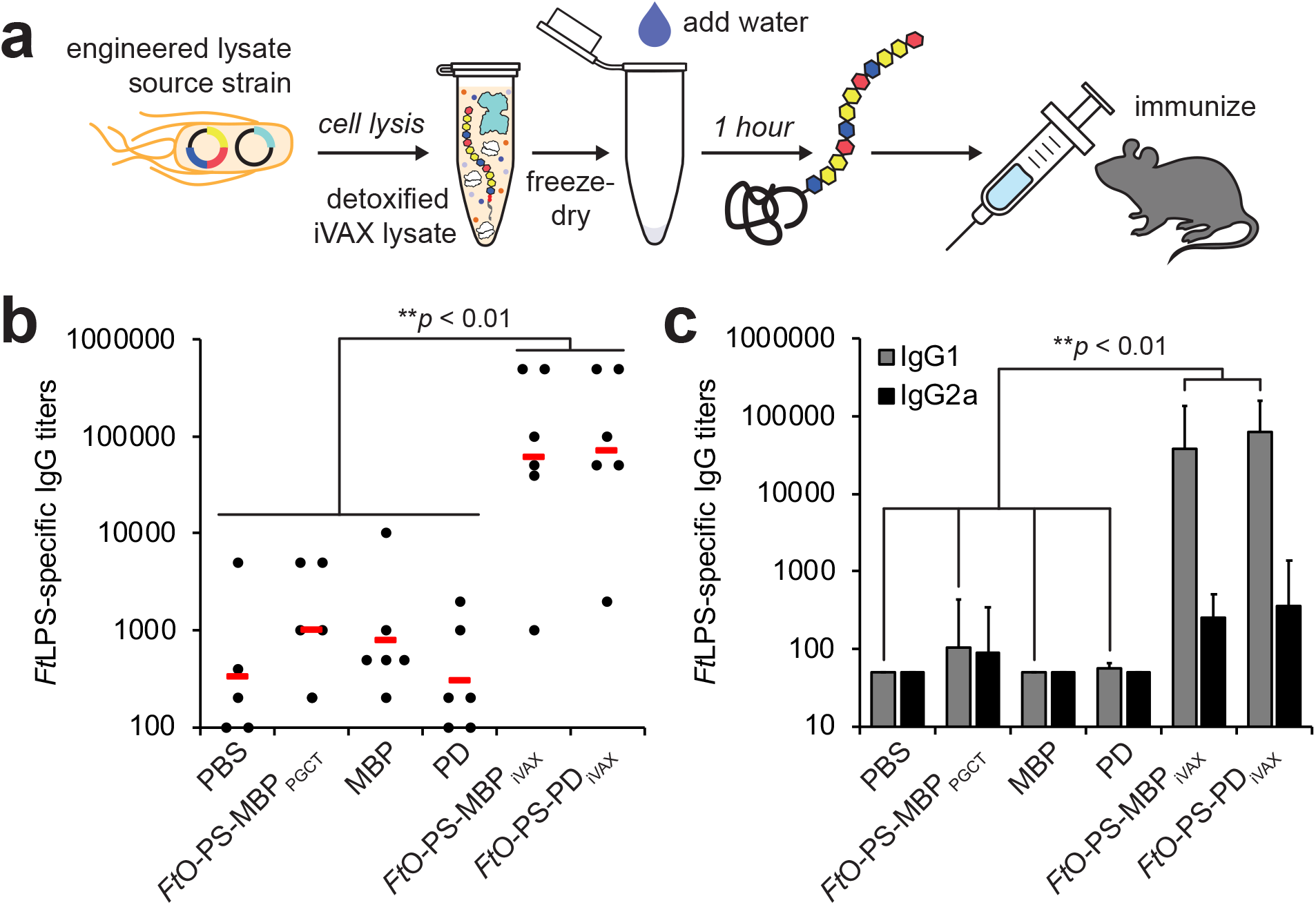
iVAX-derived bioconjugates elicit *Ft*LPS-specific antibodies in mice. (**a**) Freeze-dried iVAX reactions assembled using detoxified lysates were used to synthesize anti-*F. tularensis* bioconjugates for immunization studies. (**b**) Six groups of BALB/c mice were immunized subcutaneously with PBS or 7.5 μg of purified, cell-free synthesized aglycosylated MBP^4xDQNAT^, *Ft*O-PS-conjugated MBP^4xDQNAT^, aglycosylated PD^4xDQNAT^, or *Ft*O-PS-conjugated PD^4xDQNAT^. *Ft*O-PS-conjugated MBP^4xDQNAT^ prepared in living *E. coli* cells using PCGT was used as a positive control. Each group was composed of six mice except for the PBS control group, which was composed of five mice. Mice were boosted on days 21 and 42 with identical doses of antigen. *Ft*LPS-specific IgG titers were measured by ELISA in endpoint (day 70) serum of individual mice (black dots) with *F. tularensis* LPS immobilized as antigen. Mean titers of each group are also shown (red lines). iVAX-derived bioconjugates elicited significantly higher levels of *Ft*LPS-specific IgG compared to all other groups (\*\**p* < 0.01, Tukey-Kramer HSD). (**c**) IgG1 and IgG2a subtype titers measured by ELISA from endpoint serum revealed that iVAX-derived bioconjugates boosted production of *Ft*O-PS-specific IgG1 compared to all other groups tested (\*\**p* < 0.01, Tukey-Kramer HSD). These results indicate that iVAX bioconjugates elicited a Th2-biased immune response typical of most conjugate vaccines. Values represent means and error bars represent standard errors of *Ft*LPS-specific IgGs detected by ELISA. *See also Figure S6.*

We further characterized IgG titers by analysis of IgG1 and IgG2a subtypes and found that both iVAX-derived *Ft*O-PS-conjugated MBP^4xDQNAT^ and PD^4xDQNAT^ boosted production of IgG1 antibodies by >2 orders of magnitude relative to all control groups as well as to glycosylated MBP^4xDQNAT^ derived from PGCT (**Figure 6c**). This analysis also revealed that both iVAX-derived bioconjugates elicited a strongly Th2-biased (IgG1 >> IgG2a) response, which is characteristic of most conjugate vaccines (Bogaert et al., 2004). Taken together, these results provide evidence that the iVAX platform supplies vaccine candidates that are capable of eliciting strong, pathogen-specific humoral immune responses and recapitulate the Th2 bias that is characteristic of licensed conjugate vaccines.

## Discussion

In this work we have established iVAX, a cell-free platform for portable, on-demand production of bioconjugate vaccines. We show that iVAX reactions can be detoxified to ensure the safety of bioconjugate vaccine products, freeze-dried for cold chain-independent distribution, and re-activated for high-yielding bioconjugate production by simply adding water. As a model vaccine candidate, we show that anti-*F. tularensis* bioconjugates derived from freeze-dried, endotoxin-edited iVAX reactions elicited pathogen-specific IgG antibodies in mice as part of a Th2-biased immune response characteristic of licensed conjugate vaccines.

The iVAX platform has several important features. First, iVAX is modular, which we have demonstrated through the interchangeability of (i) carrier proteins, including those used in licensed conjugate vaccines, and (ii) bacterial O-PS antigens from *F. tularensis* subsp. *tularensis* (type A) Schu S4, ETEC *E. coli* O78, and UPEC *E. coli* O7. To our knowledge, this represents the first demonstration of oligosaccharyltransferase-mediated O-PS conjugation to authentic FDA-approved carrier proteins, likely due to historical challenges associated with the expression of licensed carriers in living *E. coli* (Figueiredo et al., 1995; Haghi et al., 2011; Stefan et al., 2011). Further expansion of the O-PS pathways used in iVAX should be possible given the commonly observed clustering of polysaccharide biosynthesis genes in the genomes of pathogenic bacteria (Raetz and Whitfield, 2002). This feature could make iVAX an attractive option for rapid, *de novo* development of bioconjugate vaccine candidates in response to a disease outbreak or against emerging drug-resistant bacteria.

Second, iVAX reactions are inexpensive, costing ~$12 mL^−1^ (**Table S1**) with the ability to synthesize ~20 μg bioconjugate mL^−1^ in one hour (**Figure S4a**). Assuming a dose size of 10 μg, our iVAX reactions can produce a vaccine dose for ~$6. For comparison, the CDC cost per dose for conjugate vaccines ranges from ~$9.50 for the *H. influenzae* vaccine ActHIB^®^ to ~$75 and ~$118 for the meningococcal vaccine Menactra^®^ and pneumococcal vaccine Prevnar 13^®^, respectively (CDC, 2019).

Third, and rather interestingly, we observed that iVAX-derived bioconjugates were significantly more effective at eliciting *Ft*LPS-specific IgGs than a bioconjugate derived from living *E. coli* cells using PGCT (**Figure 6**). One possible explanation for this increased effectiveness is the more extensive glycosylation that we observed for the *in vitro* expressed bioconjugates, with greater carbohydrate loading and decoration with higher molecular weight *Ft*O-PS species compared to their PGCT-derived counterparts. The reduced presence of high molecular weight O-PS species observed on bioconjugates produced using PGCT could be due to competition between the O-antigen polymerase Wzy and PglB *in vivo*. In contrast, our *in vitro* approach decouples O-PS synthesis, which occurs *in vivo* before lysate production, from glycosylation, which occurs *in vitro* as part of iVAX reactions. Our results are consistent with previous reports of PGCT-derived anti-*F. tularensis* bioconjugates which show that increasing the ratio of conjugated *Ft*O-PS to protein in bioconjugates yields enhanced protection against *F. tularensis* Schu S4 in a rat inhalation model of tularemia (Marshall et al., 2018). In addition, a recent study showed that a conjugate vaccine made with a high molecular weight (80 kDa) *Ft*O-PS coupled to TT conferred better protection against intranasal challenge with *F. tularensis* live vaccine strain compared to a conjugate made with a low molecular weight (25 kDa) polysaccharide (Stefanetti et al., 2019). These results as well as our own point to the fact that a deeper understanding of important immunogen design features such as glycan loading and chain length could enable the production of more effective conjugate vaccines in the future.

Importantly, by enabling portable production of bioconjugate vaccines, iVAX addresses a key gap in both cell-free and decentralized biomanufacturing technologies. Production of glycosylated products has not yet been demonstrated in cell-based decentralized biomanufacturing platforms (Crowell et al., 2018; Perez-Pinera et al., 2016) and existing cell-free platforms using *E. coli* lysates lack the ability to synthesize glycoproteins (Boles et al., 2017; Murphy et al., 2019; Pardee et al., 2016; Salehi et al., 2016). While glycosylated human erythropoietin has been produced in a cell-free biomanufacturing platform based on freeze-dried Chinese hamster ovary cell lysates (Adiga et al., 2018), this and the vast majority of other eukaryotic cell-free and cell-based systems rely on endogenous protein glycosylation machinery. As a result, these expression platforms offer little control over the glycan structures produced or the underlying glycosylation reactions, and significant optimization is often required to achieve acceptable glycosylation profiles (Adiga et al., 2018). In contrast, the iVAX platform is enabled by lysates derived from *E. coli* that lack endogenous protein glycosylation pathways, allowing for bottom-up engineering of desired glycosylation activity (Jaroentomeechai et al., 2018). The ability to engineer desired glycosylation activity in iVAX uniquely enables the production of an important class of antibacterial vaccines at the point-of-care.

In summary, iVAX provides a new approach for rapid development and portable, on-demand biomanufacturing of bioconjugate vaccines. The iVAX platform alleviates cold chain requirements, which could enhance delivery of medicines to regions with limited infrastructure and minimize vaccine losses due to spoilage. In addition, the ability to rapidly produce vaccine candidates *in vitro* provides a unique means for rapidly responding to pathogen outbreaks and emergent threats. As a result, we believe that the iVAX platform, along with an emerging set of technologies with the ability to synthesize biomedicines on-demand (Adiga et al., 2018; Boles et al., 2017; Crowell et al., 2018; Murphy et al., 2019; Pardee et al., 2016; Perez-Pinera et al., 2016; Salehi et al., 2016), has the potential to promote increased access to complex, life-saving drugs through decentralized production.

## Acknowledgements

This work was supported by Defense Threat Reduction Agency (HDTRA1-15-10052/P00001 to M.P.D. and M.C.J.), National Science Foundation (Grants # CBET 1159581 and CBET 1264701 both to M.P.D. and MCB 1413563 to M.P.D. and M.C.J.), the David and Lucile Packard Foundation, and the Dreyfus Teacher-Scholar program. J.C.S. and T.C.S. were supported by National Science Foundation Graduate Research Fellowships. T.J. was supported by a Royal Thai Government Fellowship. R.S.D. was funded, in part, by the Northwestern University Chemistry of Life Processes Summer Scholars program.

## Author Contributions

J.C.S. designed research, performed research, analyzed data, and wrote the paper. T.J. and T.M. designed research, performed research, and analyzed data. R.S.D. and K.J.H. performed research. T.C.S. aided in research design. M.C.J. and M.P.D. directed research, analyzed data, and wrote the paper.

## STAR Methods

### Contact for Reagent and Resource Sharing

For further information and request of reagents please address the lead contact Michael Jewett.

### Experimental Model and Subject Details

#### Bacteria

*E. coli* strains were grown in Luria Bertani (LB), LB-Lennox, or 2xYTP media at 30-37°C in shaking incubators, as specified in the Method Details section below. *E. coli* NEB 5-alpha (NEB) was used for plasmid cloning and purification. *E. coli* CLM24 or CLM24 Δ*lpxM* strains were used for preparing cell-free lysates. *E. coli* CLM24 was used as the chassis for expressing bioconjugates *in vivo* using PGCT. CLM24 is a glyco-optimized derivative of W3110 that carries a deletion in the gene encoding the WaaL ligase, facilitating the accumulation of preassembled glycans on undecaprenyl diphosphate (Feldman et al., 2005). CLM24 Δ*lpxM* has an endogenous acyltransferase deletion and serves as the chassis strain for production of detoxified cell-free lysates.

#### Human cell culture

HEK-Blue hTLR4 cells (Invivogen) were cultured in DMEM media high glucose/L-glutamine supplement with 10% fetal bovine serum, 50 U mL^−1^ penicillin, 50 mg mL^−1^ streptomycin, and 100 μg mL^−1^ Normacin™ in a humidified incubator at 37°C with 5% CO_2_, per manufacturer’s specifications.

#### Mice

Six-week old female BALB/c mice obtained from Harlan Sprague Dawley were randomly assigned to experimental groups. Mice were housed in groups of three to five animals and cages were changed weekly. All animals were observed daily for behavioral and physical health such as visible signs of injury and abnormal grooming habits. Mice were also observed 24 and 48 hours after each injection and no abnormal responses were reported. This study was performed at Cornell University in facilities accredited by the Association for Assessment and Accreditation of Laboratory Animal Care International. All procedures were done in accordance with Protocol 2012-0132 approved by the Cornell University Institutional Animal Care and Use Committee under relevant university, state, and federal regulations including the Animal Welfare Act, Public Health Service Policy on Humane Care and Use of Laboratory Animals, New York State Health Law Article 5, Section 504, and Cornell University Policy 1.4.

### Method Details

#### Construction of CLM24 ΔlpxM strain

*E. coli* CLM24 Δ*lpxM* was generated using the Datsenko-Wanner gene knockout method (Datsenko and Wanner, 2000). Briefly, CLM24 cells were transformed with the pKD46 plasmid encoding the λ red system. Transformants were grown to an OD_600_ of 0.5-0.7 in 25 mL LB-Lennox media (10 g L^−1^ tryptone, 5 g L^−1^ yeast extract and 5 g L^−1^ NaCl) with 50 μg mL^−1^ carbenicillin at 30°C, harvested and washed three times with 25 mL ice-cold 10% glycerol to make them electrocompetent, and resuspended in a final volume of 100 μL 10% glycerol. In parallel, a *lpxM* knockout cassette was generated by PCR amplifying the kanamycin resistance cassette from pKD4 with forward and reverse primers with homology to *lpxM*. Electrocompetent cells were transformed with 400 ng of the *lpxM* knockout cassette and plated on LB agar with 30 μg mL^−1^ kanamycin for selection of resistant colonies. Plates were grown at 37°C to cure cells of the pKD46 plasmid. Colonies that grew on kanamycin were confirmed to have acquired the knockout cassette via colony PCR and DNA sequencing. These confirmed colonies were then transformed with pCP20 to remove the kanamycin resistance gene via Flp-FRT recombination. Transformants were plated on LB agar with 50 μg mL^−1^ carbenicillin. Following selection, colonies were grown in liquid culture at 42°C to cure cells of the pCP20 plasmid. Colonies were confirmed to have lost both *lpxM* and the knockout cassette via colony PCR and DNA sequencing and confirmed to have lost both kanamycin and carbenicillin resistance via replica plating on LB agar plates with 50 μg mL^−1^ carbenicillin or kanamycin. All primers used for construction and validation of this strain are listed in **Table S2**.

All plasmids used in the study are listed in **Table S3**. Plasmids pJL1-MBP^4xDQNAT^, pJL1-PD^4xDQNAT^, pJL1-PorA^4xDQNAT^, pJL1-TTc^4xDQNAT^, pJL1-TTlight^4xDQNAT^, pJL1-CRM197^4xDQNAT^, and pJL1-TT^4xDQNAT^ were generated via PCR amplification and subsequent Gibson Assembly of a codon optimized gene construct purchased from IDT with a C-terminal 4xDQNAT-6xHis tag (Fisher et al., 2011) between the *NdeI* and *SalI* restriction sites in the pJL1 vector. Plasmid pJL1-EPA^DNNNS-DQNRT^ was constructed using the same approach, but without the addition of a C-terminal 4xDQNAT-6xHis tag. Plasmids pTrc99s-ssDsbA-MBP^4xDQNAT^, pTrc99s-ssDsbA-PD^4xDQNAT^, pTrc99s-ssDsbA-PorA^4xDQNAT^, pTrc99s-ssDsbA-TTc^4xDQNAT^, pTrc99s-ssDsbA-TTlight^4xDQNAT^, and pTrc99s-ssDsbA-EPA^DNNNS-DQNRT^ were created via PCR amplification of each carrier protein gene and insertion into the pTrc99s vector between the *NcoI* and *HindIII* restriction sites via Gibson Assembly. Plasmid pSF-*Cj*PglB-LpxE was constructed using a similar approach, but via insertion of the *lpxE* gene from pE (Needham et al., 2013) between the *NdeI* and *NsiI* restriction sites in the pSF vector. Inserts were amplified via PCR using Phusion® High-Fidelity DNA polymerase (NEB) with forward and reverse primers designed using the NEBuilder® Assembly Tool (nebuilder.neb.com) and purchased from IDT. The pJL1 vector (Addgene 69496) was digested using restriction enzymes NdeI and SalI-HF® (NEB). The pSF vector was digested using restriction enzymes NdeI and NotI (NEB). PCR products were gel extracted using an EZNA Gel Extraction Kit (Omega Bio-Tek), mixed with Gibson assembly reagents and incubated at 50°C for 1 hour. Plasmid DNA from the Gibson assembly reactions were transformed into *E. coli* NEB 5-alpha cells and circularized constructs were selected using kanamycin at 50 μg ml^−1^ (Sigma). Sequence-verified clones were purified using an EZNA Plasmid Midi Kit (Omega Bio-Tek) for use in CFPS and iVAX reactions.

#### Cell-free lysate preparation

*E. coli* CLM24 source strains were grown in 2xYTP media (10 g/L yeast extract, 16 g/L tryptone, 5 g/L NaCl, 7 g/L K_2_HPO_4_, 3 g/L KH_2_PO_4_, pH 7.2) in shake flasks (1 L scale) or a Sartorius Stedim BIOSTAT Cplus bioreactor (10 L scale) at 37°C. Protein synthesis yields and glycosylation activity were reproducible across different batches of lysate at both small and large scale. To generate *Cj*PglB-enriched lysate, CLM24 cells carrying plasmid pSF-*Cj*PglB (Ollis et al., 2015) was used as the source strain. To generate *Ft*O-PS-enriched lysates, CLM24 carrying plasmid pGAB2 (Cuccui et al., 2013) was used as the source strain. To generate one-pot lysates containing both *Cj*PglB and *Ft*O-PS, *Ec*O78-PS, or *Ec*O7-PS, CLM24 carrying pSF-*Cj*PglB and one of the following bacterial O-PS biosynthetic pathway plasmids was used as the source strain: pGAB2 (*Ft*O-PS), pMW07-O78 (*Ec*O78-PS), and pJHCV32 (*Ec*O7-PS). *Cj*PglB expression was induced at an OD_600_ of 0.8-1.0 with 0.02% (w/v) L-arabinose and cultures were moved to 30°C. Cells were grown to a final OD_600_ of ~3.0, at which point cells were pelleted by centrifugation at 5,000xg for 15 min at 4°C. Cell pellets were then washed three times with cold S30 buffer (10 mM Tris-acetate pH 8.2, 14 mM magnesium acetate, 60 mM potassium acetate) and pelleted at 5000xg for 10 min at 4°C. After the final wash, cells were pelleted at 7000xg for 10 min at 4°C, weighed, flash frozen in liquid nitrogen, and stored at −80°C. To make cell lysate, cell pellets were resuspended to homogeneity in 1 mL of S30 buffer per 1 g of wet cell mass. Cells were disrupted via a single passage through an Avestin EmulsiFlex-B15 (1 L scale) or EmulsiFlex-C3 (10 L scale) high-pressure homogenizer at 20,000-25,000 psi. The lysate was then centrifuged twice at 30,000×g for 30 min to remove cell debris. Supernatant was transferred to clean microcentrifuge tubes and incubated at 37°C with shaking at 250 rpm for 60 min. Following centrifugation (15,000xg) for 15 min at 4°C, supernatant was collected, aliquoted, flash-frozen in liquid nitrogen, and stored at −80°C. S30 lysate was active for about 3 freeze-thaw cycles and contained ~40 g/L total protein as measured by Bradford assay.

#### Cell-free protein synthesis

CFPS reactions were carried out in 1.5 mL microcentrifuge tubes (15 μL scale), 15 mL conical tubes (1 mL scale), or 50 mL conical tubes (5 mL scale) with a modified PANOx-SP system (Jewett and Swartz, 2004). The CFPS reaction mixture consists of the following components: 1.2 mM ATP; 0.85 mM each of GTP, UTP, and CTP; 34.0 μg mL^−1^ L-5-formyl-5, 6, 7, 8-tetrahydrofolic acid (folinic acid); 170.0 μg mL^−1^ of *E. coli* tRNA mixture; 130 mM potassium glutamate; 10 mM ammonium glutamate; 12 mM magnesium glutamate; 2 mM each of 20 amino acids; 0.4 mM nicotinamide adenine dinucleotide (NAD); 0.27 mM coenzyme-A (CoA); 1.5 mM spermidine; 1 mM putrescine; 4 mM sodium oxalate; 33 mM phosphoenolpyruvate (PEP); 57 mM HEPES; 13.3 μg mL^−1^ plasmid; and 27% v/v of cell lysate. For reaction volumes ≥1 mL, plasmid was added at 6.67 μg mL^−1^, as this lower plasmid concentration conserved reagents with no effect on protein synthesis yields or kinetics. For expression of PorA, reactions were supplemented with nanodiscs at 1 μg mL^−1^, which were prepared as previously described (Schoborg et al., 2017) or purchased (Cube Biotech). For expression of CRM197^4xDQNAT^, CFPS was carried out at 25°C for 20 hours, unless otherwise noted. For all other carrier proteins, CFPS was run at 30°C for 20 hours, unless otherwise noted.

For expression of TT^4xDQNAT^, which contains intermolecular disulfide bonds, CFPS was carried out under oxidizing conditions. For oxidizing conditions, lysate was pre-conditioned with 750 μM iodoacetamide at room temperature for 30 min to covalently bind free sulfhydryls (-SH) and the reaction mix was supplemented with 200 mM glutathione at a 4:1 ratio of oxidized and reduced forms and 10 μM recombinant *E. coli* DsbC (Knapp et al., 2007).

#### <u>I</u>n vitro bioconjugate <u>va</u>ccine e<u>x</u>pression (iVAX)

For *in vitro* expression and glycosylation of carrier proteins in crude lysates, a two-phase scheme was implemented. In the first phase, CFPS was carried out for 15 min at 25-30°C as described above. In the second phase, protein glycosylation was initiated by the addition of MnCl_2_ and DDM at a final concentration of 25 mM and 0.1% (w/v), respectively, and allowed to proceed at 30°C for a total reaction time of 1 hour. Protein synthesis yields and glycosylation activity were reproducible across biological replicates of iVAX reactions at both small and large scale. Reactions were then centrifuged at 20,000xg for 10 min to remove insoluble or aggregated protein products and the supernatant was analyzed by SDS-PAGE and Western blotting.

Purification of aglycosylated and glycosylated carriers from iVAX reactions was carried out using Ni-NTA agarose (Qiagen) according to manufacturer’s protocols. Briefly, 0.5 mL Ni-NTA agarose per 1 mL cell-free reaction mixture was equilibrated in Buffer 1 (300 mM NaCl 50 mM NaH_2_PO_4_) with 10 mM imidazole. Soluble fractions from iVAX reactions were loaded on Ni-NTA agarose and incubated at 4°C for 2-4 hours to bind 6xHis-tagged protein. Following incubation, the cell-free reaction/agarose mixture was loaded onto a polypropylene column (BioRad) and washed twice with 6 column volumes of Buffer 1 with 20 mM imidazole. Protein was eluted in 4 fractions, each with 0.3 mL Buffer 1 with 300 mM imidazole per mL of cell-free reaction mixture. All buffers were used and stored at 4°C. Protein was stored at a final concentration of 1-2 mg mL^−1^ in sterile 1xPBS (137 mM NaCl, 2.7 mM KCl, 10 mM Na_2_HPO_4_, 1.8 mM KH_2_PO_4_, pH 7.4) at 4°C.

#### Expression of bioconjugates in vivo using <u>p</u>rotein <u>g</u>lycan <u>c</u>oupling <u>t</u>echnology (PGCT)

Plasmids encoding bioconjugate carrier protein genes preceded by the DsbA leader sequence for translocation to the periplasm were transformed into CLM24 cells carrying pGAB2 and pSF-*Cj*PglB. CLM24 carrying only pGAB2 was used as a negative control. Transformed cells were grown in 5 mL LB media (10 g L^−1^ yeast extract, 5 g L^−1^ tryptone, 5 g L^−1^ NaCl) overnight at 37°C. The next day, cells were subcultured into 100 mL LB and allowed to grow at 37°C for 6 hours after which the culture was supplemented with 0.2% arabinose and 0.5 mM IPTG to induce expression of *Cj*PglB and the bioconjugate carrier protein, respectively. Protein expression was then carried out for 16 hours at 30°C, at which point cells were harvested. Cell pellets were resuspended in 1 mL sterile PBS (137 mM NaCl, 2.7 mM KCl, 10 mM Na_2_HPO_4_, 1.8 mM KH_2_PO_4_, pH 7.4) and lysed using a Q125 Sonicator (Qsonica, Newtown, CT) at 40% amplitude in cycles of 10 sec on/10 sec off for a total of 5 min. Soluble fractions were isolated following centrifugation at 15,000 rpm for 30 min at 4°C. Protein was purified from soluble fractions using Ni-NTA spin columns (Qiagen), following the manufacturer’s protocol.

#### Western blot analysis

Samples were run on 4-12% Bis-Tris SDS-PAGE gels (Invitrogen). Following electrophoretic separation, proteins were transferred from gels onto Immobilon-P polyvinylidene difluoride (PVDF) membranes (0.45 μm) according to the manufacturer’s protocol. Membranes were washed with PBS (80 g L^−1^ NaCl, 0.2 g L^−1^ KCl, 1.44 g L^−1^ Na_2_HPO_4_, 0.24 g L^−1^ KH_2_PO_4_, pH 7.4) followed by incubation for 1 hour in Odyssey® Blocking Buffer (LI-COR). After blocking, membranes were washed 6 times with PBST (80 g L^−1^ NaCl, 0.2 g L^−1^ KCl, 1.44 g L^−1^ Na_2_HPO_4_, 0.24 g L^−1^ KH_2_PO_4_, 1 mL L^−1^ Tween-20, pH 7.4) with a 5 min incubation between each wash. For iVAX samples, membranes were probed with both an anti-6xHis tag antibody and an anti-O-PS antibody or antisera specific to the O antigen of interest, if commercially available. Probing of membranes was performed for at least 1 hour with shaking at room temperature, after which membranes were washed with PBST in the same manner as described above and probed with fluorescently labeled secondary antibodies. Membranes were imaged using an Odyssey® Fc imaging system (LI-COR). CRM197 and TT production were compared to commercial DT and TT standards (Sigma) and orthogonally detected by an identical SDS-PAGE procedure followed by Western blot analysis with a polyclonal antibody that recognizes diphtheria or tetanus toxin, respectively. All antibodies and dilutions used are listed in **Table S4**.

#### TLR4 activation assay

HEK-Blue hTLR4 cells (Invivogen) were maintained in DMEM media, high glucose/L-glutamine supplement with 10% fetal bovine serum, 50 U mL^−1^ penicillin, 50 mg mL^−1^ streptomycin, and 100 μg mL^−1^ Normacin™ at 37°C in a humidified incubator containing 5% CO_2_. After reaching ~50-80% confluency, cells were plated into 96-well plates at a density of 1.4 × 10^5^ cells per mL in HEK-Blue detection media (Invivogen). Antigens were added at the following concentrations: 100 ng μL^−1^ purified protein; and 100 ng μL^−1^ total protein in lysate. Purified *E. coli* O55:B5 LPS (Sigma-Aldrich) and detoxified *E. coli* O55:B5 (Sigma-Aldrich) were added at 1.0 ng mL^−1^ and served as positive and negative controls, respectively. Plates were incubated at 37°C, 5% CO_2_ for 10–16 h before measuring absorbance at 620 nm. Statistical significance was determined using paired *t-*tests.

#### Mouse immunization

Six groups of six-week old female BALB/c mice (Harlan Sprague Dawley) were injected subcutaneously with 100 μL PBS (pH 7.4) alone or containing purified aglycosylated MBP, *Ft*O-PS-conjugated MBP, aglycosylated PD, or *Ft*O-PS-conjugated PD, as previously described (Chen et al., 2010). Groups were composed of six mice except for the PBS control group, which was composed of five mice. The amount of antigen in each preparation was normalized to 7.5 μg to ensure that an equivalent amount of aglycosylated protein or bioconjugate was administered in each case. Purified protein groups formulated in PBS were mixed with an equal volume of incomplete Freund’s Adjuvant (Sigma-Aldrich) before injection. Prior to immunization, material for each group (5 μL) was streaked on LB agar plates and grown overnight at 37°C to confirm sterility and endotoxin activity was measured by TLR4 activation assay. Each group of mice was boosted with an identical dosage of antigen 21 days and 42 days after the initial immunization. Blood was obtained on day −1, 21, 35, 49, and 63 via submandibular collection and at study termination on day 70 via cardiac puncture. Mice were observed 24 and 48 hours after each injection for changes in behavior and physical health and no abnormal responses were reported. This study and all procedures were done in accordance with Protocol 2012-0132 approved by the Cornell University Institutional Animal Care and Use Committee.

#### Enzyme-linked immunosorbent assay

*F. tularensis* LPS-specific antibodies elicited by immunized mice were measured via indirect ELISA using a modification of a previously described protocol (Chen et al., 2010). Briefly, sera were isolated from the collected blood draws after centrifugation at 5000xg for 10 min and stored at −20 °C; 96-well plates (Maxisorp; Nunc Nalgene) were coated with *F. tularensis* LPS (BEI resources) at a concentration of 5 μg mL^−1^ in PBS and incubated overnight at 4°C. The next day, plates were washed three times with PBST (PBS, 0.05% Tween-20, 0.3% BSA) and blocked overnight at 4°C with 5% nonfat dry milk (Carnation) in PBS. Samples were serially diluted by a factor of two in triplicate between 1:100 and 1:12,800,000 in blocking buffer and added to the plate for 2 hours at 37°C. Plates were washed three times with PBST and incubated for 1 hour at 37°C in the presence of one of the following HRP-conjugated antibodies (all from Abcam and used at 1:25,000 dilution): goat anti-mouse IgG, anti-mouse IgG1, and anti-mouse IgG2a. After three additional washes with PBST, 3,3′-5,5′-tetramethylbenzidine substrate (1-Step Ultra TMB-ELISA; Thermo-Fisher) was added, and the plate was incubated at room temperature in the dark for 30 min. The reaction was halted with 2 M H_2_SO_4_, and absorbance was quantified via microplate spectrophotometer (Tecan) at a wavelength of 450 nm. Serum antibody titers were determined by measuring the lowest dilution that resulted in signal 3 SDs above no serum background controls. Statistical significance was determined in RStudio 1.1.463 using one-way ANOVA and the Tukey–Kramer *post hoc* honest significant difference test.

### Quantification and Statistical Analysis

#### Quantification of cell-free protein synthesis yields

To quantify the amount of protein synthesized in iVAX reactions, two approaches were used. Fluorescence units of sfGFP were converted to concentrations using a previously reported standard curve (Hong et al., 2014). Yields of all other proteins were assessed via the addition of 10 μM L-^14^C-leucine (11.1 GBq mmol^−1^, PerkinElmer) to the CFPS mixture to yield trichloroacetic acid-precipitable radioactivity that was measured using a liquid scintillation counter as described previously (Kim and Swartz, 2001).

#### Statistical analysis

Statistical parameters including the definitions and values of *n*, *p*-values, standard deviations, and standard errors are reported in the Figures and corresponding Figure Legends. Analytical techniques are described in the corresponding Method Details section.

## Data and Software Availability

All plasmid constructs used in this study including complete DNA sequences are deposited on Addgene (constructs 128389-128404).

## Supplemental Information Titles and Legends

**Figure S1. *In vitro* synthesis of licensed conjugate vaccine carrier proteins is possible over a range of temperatures and can be readily optimized, Related to Figure 2.** (**a**) With the exception of CRM197, all carriers expressed with similar soluble yields at 25°C, 30°C, and 37°C, as measured by ^14^C-leucine incorporation. Values represent means and error bars represent standard deviations of biological replicates (*n* = 3). (**b**) Soluble expression of PorA was improved through the addition of lipid nanodiscs to the reaction. (**c**) Expression of full-length TT was enhanced by (i) performing *in vitro* protein synthesis in oxidizing conditions to improve assembly of the disulfide-bonded heavy and light chains into full-length TT and (ii) allowing reactions to run for only 2 h to minimize protease degradation. (**d**) CRM197 and (**e**) TT produced in CFPS reactions are detected with α-DT and α-TT antibodies, respectively, and are comparable in size to commercially available purified DT and TT protein standards (50 ng standard loaded). Images are representative of at least three biological replicates. Dashed line indicates samples are from the same blot with the same exposure. Molecular weight ladders are shown at the left of each image.

**Figure S2. Glycosylation in iVAX reactions occurs in 1 h over a range of temperatures, Related to Figure 3.** Kinetics of *Ft*O-PS glycosylation at 30°C (**left**), 37°C, 25°C, and room temperature (~21°C) (**right**) are comparable and show that protein synthesis and glycosylation occur in the first hour of the iVAX reaction. These results demonstrate that the iVAX platform can synthesize bioconjugates over a range of permissible temperatures. Top panels show signal from probing with anti-hexa-histidine antibody (αHis) to detect the carrier protein, middle panels show signal from probing with commercial anti-*Ft*O-PS antibody (α*Ft*O-PS), and bottom panels show αHis and α*Ft*O-PS signals merged. Images are representative of at least three biological replicates. Molecular weight ladders are shown at the left of each image.

**Figure S3. Production of bioconjugates against *F. tularensis* using PGCT in living *E. coli*, Related to Figure 4.** (**a**) Bioconjugates were produced via PGCT in CLM24 cells expressing *Cj*PglB, the biosynthetic pathway for *Ft*O-PS, and a panel of immunostimulatory carriers including those used in licensed vaccines. (**b**) We observed low expression of PorA, a membrane protein, as well as reduced glycan loading and conjugation of high molecular weight *Ft*O-PS species in all carriers compared to iVAX-derived samples. Top panels show signal from probing with anti-hexa-histidine antibody (αHis) to detect the carrier protein, middle panels show signal from probing with commercial anti-*Ft*O-PS antibody (α*Ft*O-PS), and bottom panels show αHis and α*Ft*O-PS signals merged. Images are representative of at least three biological replicates. Molecular weight ladders are shown at the left of each image.

**Figure S4. The iVAX platform is modular and can be used to synthesize clinically relevant yields of diverse bioconjugates, Related to Figure 4.** (**a**) Protein synthesis and glycosylation with *Ft*O-PS were measured in iVAX reactions producing MBP^4xDQNAT^ and PD^4xDQNAT^. After ~1 h, reactions produced ~40 μg mL^−1^ protein, as measured via ^14^C-leucine incorporation, of which ~20 μg mL^−1^ was glycosylated with *Ft*O-PS, as determined by densitometry. Values represent means and error bars represent standard errors of biological replicates (*n* = 2). To demonstrate modularity, iVAX lysates were prepared from cells expressing *Cj*PglB and biosynthetic pathways for either (**b**) the *E. coli* O78 antigen or (**c**) the *E. coli* O7 antigen and used to synthesize PD^4xDQNAT^ (**left**) or sfGFP^217-DQNAT^ (**right**) bioconjugates. The structure and composition of the repeating monomer unit for each antigen is shown. Both polysaccharide antigens are compositionally and, in the case of the O7 antigen, structurally distinct compared to the *F. tularensis* O antigen. Blots show signal from probing with anti-hexa-histidine antibody (αHis) to detect the carrier protein. If a commercial anti-O-PS serum or antibody was available, it was used to confirm the identity of the conjugated O antigen (α-*Ec*O78 blots, panel **b**). Asterisk denotes bands resulting from non-specific serum antibody binding. Images are representative of at least three biological replicates. Dashed lines indicate samples are from the same blot with the same exposure. Molecular weight ladders are shown at the left of each image.

**Figure S5. Detoxified iVAX lysates synthesize bioconjugates and both lysate production and freeze-dried reactions scale reproducibly, Related to Figure 5.** (**a**) iVAX lysates containing *Cj*PglB and *Ft*O-PS were prepared from wild-type CLM24, CLM24 Δ*lpxM*, or CLM24 Δ*lpxM* cells expressing *Ft*LpxE. Nearly identical sfGFP^217-DQNAT^ glycosylation was observed for each of the lysates derived from the engineered strains. (**b**) To generate material for immunizations, fermentations to produce endotoxin-edited iVAX lysates were scaled from 0.5 L to 10 L. We observed similar levels of sfGFP^217-DQNAT^ glycosylation for lysates derived from 0.5 L and 10 L cultures, and across different batches of lysate produced from 10 L fermentations. (**c**) For immunizations, we prepared two lots of *Ft*O-PS-conjugated MBP^4xDQNAT^ and PD^4xDQNAT^ from 5 mL freeze-dried iVAX reactions. We observed similar levels of purified protein (~200 μg) and *Ft*O-PS modification (>50%, measured by densitometry) across both carriers and lots of material. Top panels show signal from probing with anti-hexa-histidine antibody (αHis) to detect the carrier protein, middle panels show signal from probing with commercial anti-*Ft*O-PS antibody (α*Ft*O-PS), and bottom panels show αHis and α*Ft*O-PS signals merged. Images are representative of at least three biological replicates. Molecular weight ladders are shown at left.

**Figure S6. FtLPS-specific antibody titers in vaccinated mice over time, Related to Figure 6.** Six groups of BALB/c mice were immunized subcutaneously with PBS or 7.5 μg of purified, cell-free synthesized aglycosylated MBP^4xDQNAT^, *Ft*O-PS-conjugated MBP^4xDQNAT^, aglycosylated PD^4xDQNAT^, or *Ft*O-PS-conjugated PD^4xDQNAT^. *Ft*O-PS-conjugated MBP^4xDQNAT^ prepared in living *E. coli* cells using PCGT was used as a positive control. Each group was composed of six mice except for the PBS control group, which was composed of five mice. Mice were boosted on days 21 and 42 with identical doses of antigen. *Ft*LPS-specific IgG titers were measured by ELISA in serum collected on day −1, 35, 49, 63, and 70 following initial immunization. iVAX-derived bioconjugates elicited significantly higher levels of *Ft*LPS-specific IgG compared to compared to the PBS control group in serum collected on day 35, 49, and 70 of the study (\*\**p* < 0.01, Tukey-Kramer HSD). Values represent means and error bars represent standard errors of *Ft*LPS-specific IgGs detected by ELISA.

**Table S1. Cost analysis for iVAX reactions, Related to STAR Methods.** The total cost to assemble iVAX reactions is calculated below. A 1 mL iVAX reaction produces two 10 μg vaccine doses and can be assembled for $11.75. In the table, amino acid cost accounts for 2 mM each of the 20 canonical amino acids purchased individually from Sigma. Lysate cost is based on a single employee making 50 mL lysate from a 10 L fermentation, assuming 30 lysate batches per year and a 5-year equipment lifetime. Component source is also included in the table if it is available to purchase directly from a supplier. Homemade components cannot be purchased directly and must be prepared according to procedures described in the Methods section.

**Table S2. Primers used to generate CLM24 Δ*lpxM*, Related to STAR Methods.** Primers used to construct and verify the CLM24 Δ*lpxM* strain are listed below. KO primers were used for amplification of the kanamycin resistance cassette from pKD4 with homology to *lpxM*. Seq primers were used for colony PCRs and sequencing confirmation of knockout strains.

**Table S3. Plasmids used in this study, Related to STAR Methods.**

**Table S4. Antibodies and antisera used in this study, Related to STAR Methods.**

