## Supplemental Information for "On-demand, cell-free biomanufacturing of conjugate vaccines at the point-of-care"

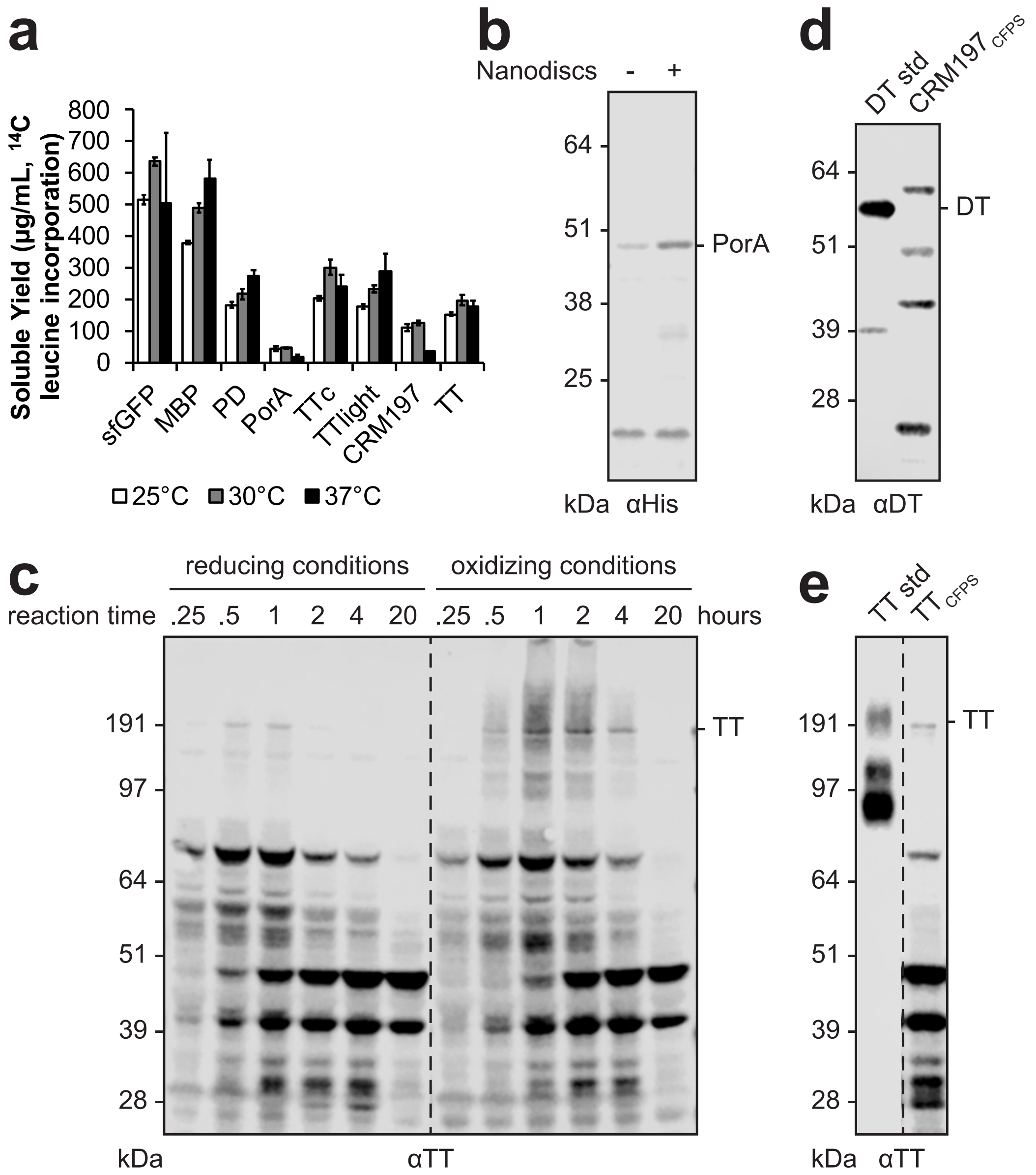


**Figure S1. *In vitro* synthesis of licensed conjugate vaccine carrier proteins is possible over a range of temperatures and can be readily optimized, Related to Figure 2.** (**a**) With the exception of CRM197, all carriers expressed with similar soluble yields at 25°C, 30°C, and 37°C, as measured by $^{14}$C-leucine incorporation. Values represent means and error bars represent standard deviations of biological replicates (*n* = 3). (**b**) Soluble expression of PorA was improved through the addition of lipid nanodiscs to the reaction. (**c**) Expression of full-length TT was enhanced by (i) performing *in vitro* protein synthesis in oxidizing conditions to improve assembly of the disulfide-bonded heavy and light chains into full-length TT and (ii) allowing reactions to run for only 2 h to minimize protease degradation. (**d**) CRM197 and (**e**) TT produced in CFPS reactions are detected with α-DT and α-TT antibodies, respectively, and are comparable in size to commercially available purified DT and TT protein standards (50 ng standard loaded). Images are representative of at least three biological replicates. Dashed line indicates samples are from the same blot with the same exposure. Molecular weight ladders are shown at the left of each image.

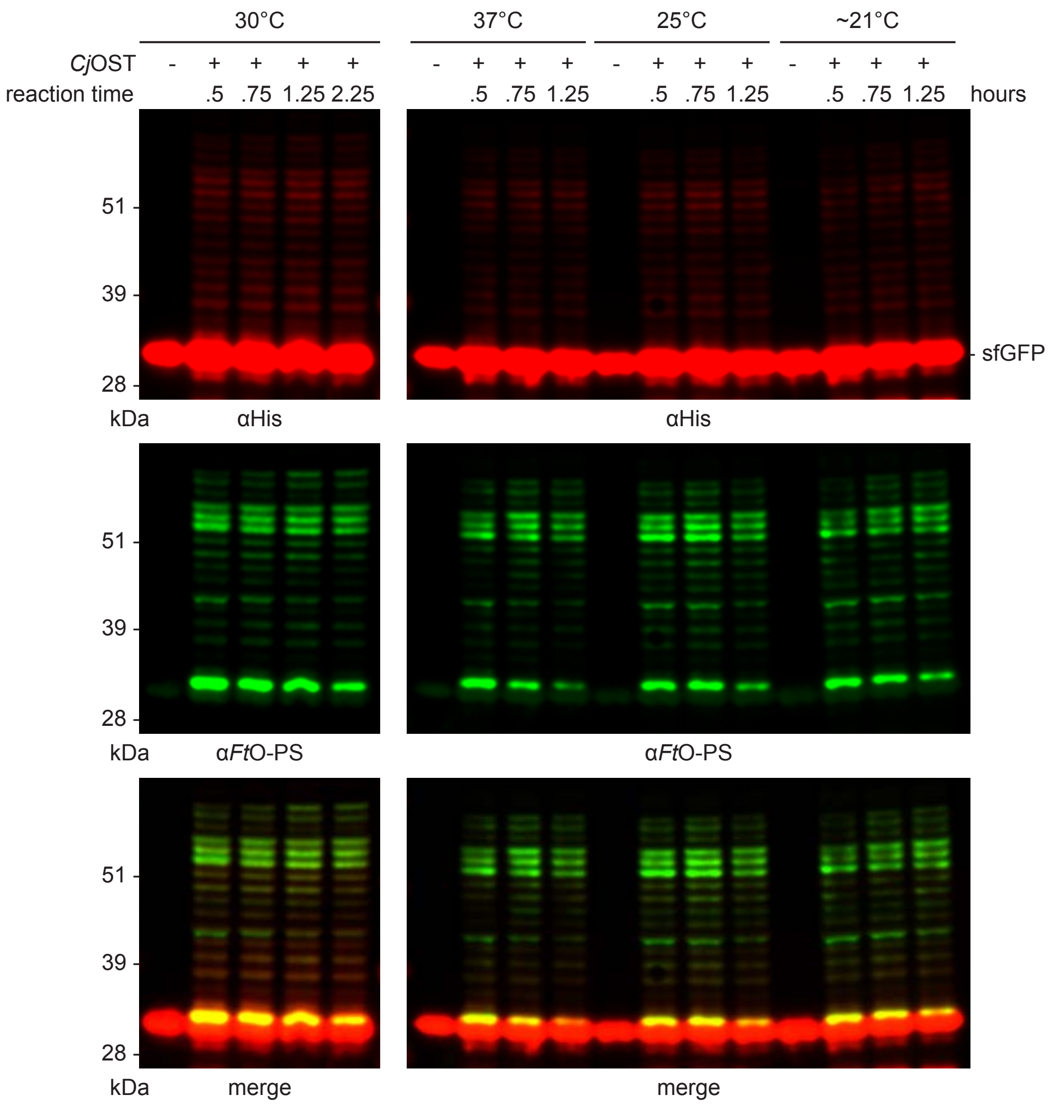


**Figure S2. Glycosylation in iVAX reactions occurs in 1 h over a range of temperatures, Related to Figure 3.** Kinetics of *Ft*O-PS glycosylation at 30°C (**left**), 37°C, 25°C, and room temperature (~21°C) (**right**) are comparable and show that protein synthesis and glycosylation occur in the first hour of the iVAX reaction. These results demonstrate that the iVAX platform can synthesize bioconjugates over a range of permissible temperatures. Top panels show signal from probing with anti-hexa-histidine antibody (αHis) to detect the carrier protein, middle panels show signal from probing with commercial anti-*Ft*O-PS antibody (α*Ft*O-PS), and bottom panels show αHis and α*Ft*O-PS signals merged. Images are representative of at least three biological replicates. Molecular weight ladders are shown at the left of each image.

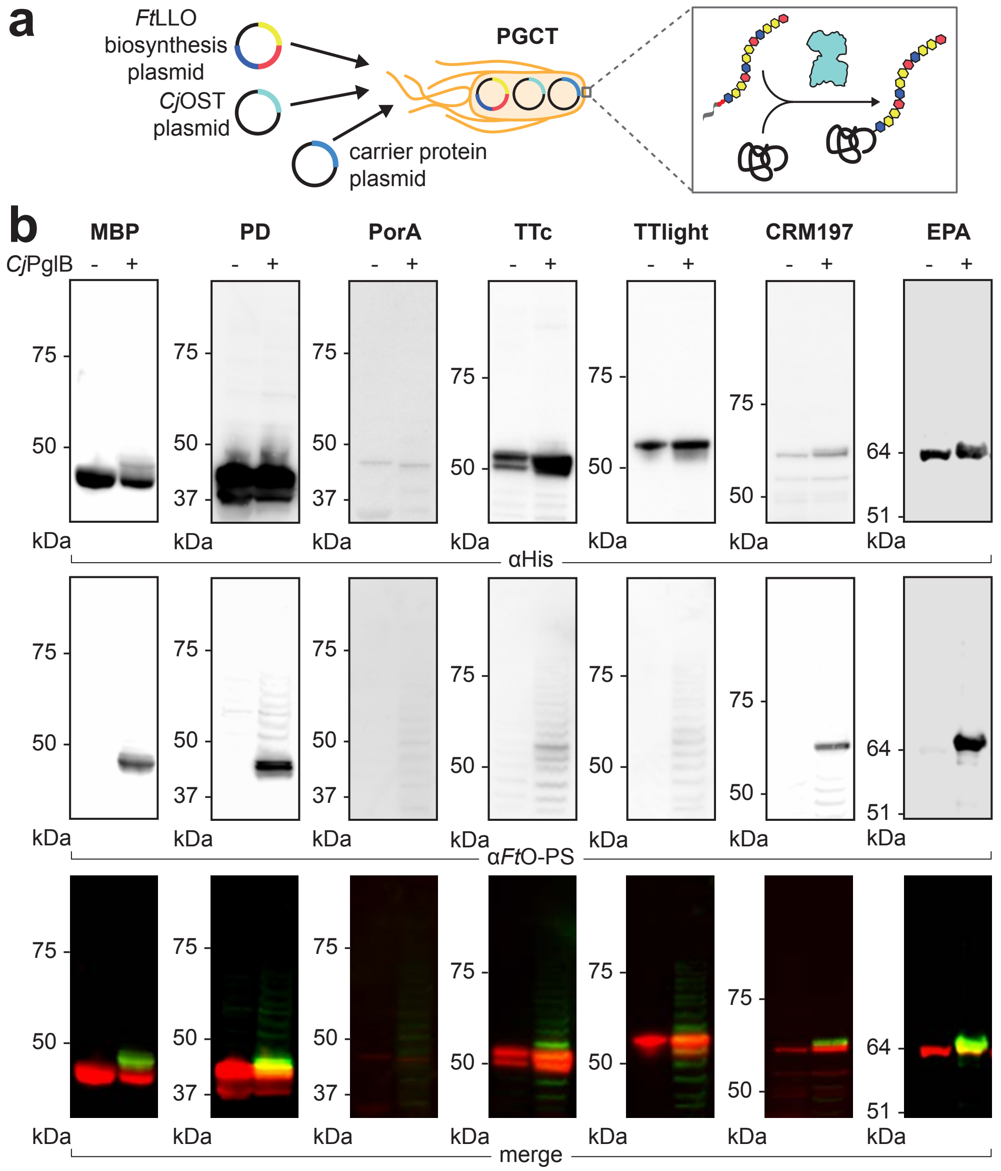


**Figure S3. Production of bioconjugates against *F. tularensis* using PGCT in living *E. coli*, Related to Figure 4.** (**a**) Bioconjugates were produced via PGCT in CLM24 cells expressing *Cj*PglB, the biosynthetic pathway for *Ft*O-PS, and a panel of immunostimulatory carriers including those used in licensed vaccines. (**b**) We observed low expression of PorA, a membrane protein, as well as reduced glycan loading and conjugation of high molecular weight *Ft*O-PS species in all carriers compared to iVAX-derived samples. Top panels show signal from probing with anti-hexa-histidine antibody (αHis) to detect the carrier protein, middle panels show signal from probing with commercial anti-*Ft*O-PS antibody (α*Ft*O-PS), and bottom panels show αHis and α*Ft*O-PS signals merged. Images are representative of at least three biological replicates. Molecular weight ladders are shown at the left of each image.

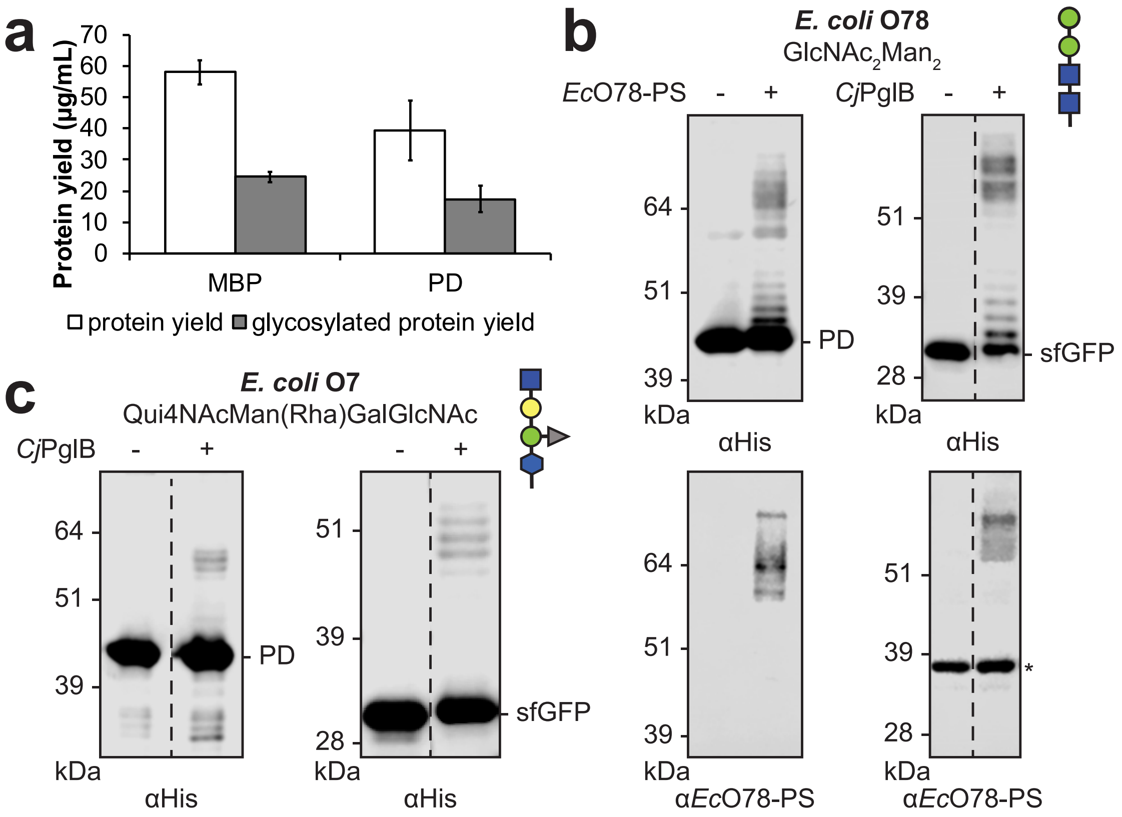


**Figure S4. The iVAX platform is modular and can be used to synthesize clinically relevant yields of diverse bioconjugates, Related to Figure 4.** (**a**) Protein synthesis and glycosylation with *Ft*O-PS were measured in iVAX reactions producing MBP$^{4xDQNAT}$ and PD$^{4xDQNAT}$. After ~1 h, reactions produced ~40 µg mL$^{-1}$ protein, as measured via $^{14}$C-leucine incorporation, of which ~20 µg mL$^{-1}$ was glycosylated with *Ft*O-PS, as determined by densitometry. Values represent means and error bars represent standard errors of biological replicates (*n* = 2). To demonstrate modularity, iVAX lysates were prepared from cells expressing *Cj*PglB and biosynthetic pathways for either (**b**) the *E. coli* O78 antigen or (**c**) the *E. coli* O7 antigen and used to synthesize PD$^{4xDQNAT}$ (**left**) or sfGFP$^{217\text{-}DQNAT}$ (**right**) bioconjugates. The structure and composition of the repeating monomer unit for each antigen is shown. Both polysaccharide antigens are compositionally and, in the case of the O7 antigen, structurally distinct compared to the *F. tularensis* O antigen. Blots show signal from probing with anti-hexa-histidine antibody (αHis) to detect the carrier protein. If a commercial anti-O-PS serum or antibody was available, it was used to confirm the identity of the conjugated O antigen (α-*Ec*O78 blots, panel **b**). Asterisk denotes bands resulting from non-specific serum antibody binding. Images are representative of at least three biological replicates. Dashed lines indicate samples are from the same blot with the same exposure. Molecular weight ladders are shown at the left of each image.

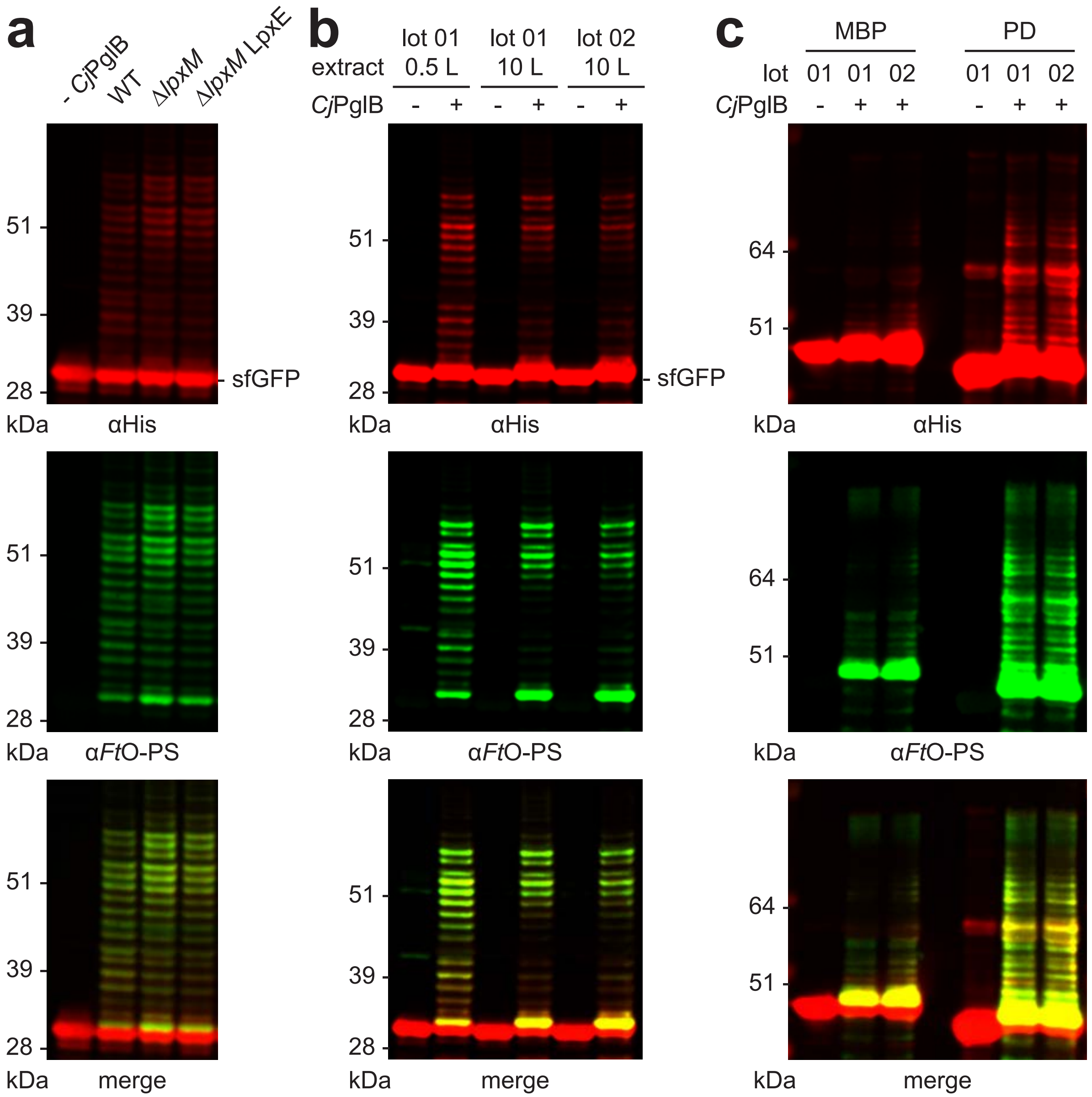


**Figure S5. Detoxified iVAX lysates synthesize bioconjugates and both lysate production and freeze-dried reactions scale reproducibly, Related to Figure 5.** (**a**) iVAX lysates containing *Cj*PglB and *Ft*O-PS were prepared from wild-type CLM24, CLM24 Δ*lpxM*, or CLM24 Δ*lpxM* cells expressing *Ft*LpxE. Nearly identical sfGFP$^{217\text{-DQNAT}}$ glycosylation was observed for each of the lysates derived from the engineered strains. (**b**) To generate material for immunizations, fermentations to produce endotoxin-edited iVAX lysates were scaled from 0.5 L to 10 L. We observed similar levels of sfGFP$^{217\text{-DQNAT}}$ glycosylation for lysates derived from 0.5 L and 10 L cultures, and across different batches of lysate produced from 10 L fermentations. (**c**) For immunizations, we prepared two lots of *Ft*O-PS-conjugated MBP$^{4\text{xDQNAT}}$ and PD$^{4\text{xDQNAT}}$ from 5 mL freeze-dried iVAX reactions. We observed similar levels of purified protein (~200 μg) and *Ft*O-PS modification (>50%, measured by densitometry) across both carriers and lots of material. Top panels show signal from probing with anti-hexa-histidine antibody (αHis) to detect the carrier protein, middle panels show signal from probing with commercial anti-*Ft*O-PS antibody (α*Ft*O-PS), and bottom panels show αHis and α*Ft*O-PS signals merged. Images are representative of at least three biological replicates. Molecular weight ladders are shown at left.

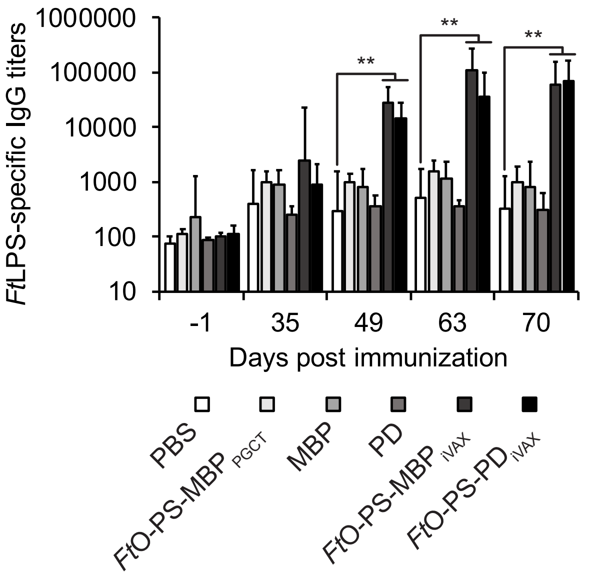


**Figure S6. *Ft*LPS-specific antibody titers in vaccinated mice over time, Related to Figure 6.** Six groups of BALB/c mice were immunized subcutaneously with PBS or 7.5 μg of purified, cell-free synthesized aglycosylated $MBP^{4xDQNAT}$, *Ft*O-PS-conjugated $MBP^{4xDQNAT}$, aglycosylated $PD^{4xDQNAT}$, or *Ft*O-PS-conjugated $PD^{4xDQNAT}$. *Ft*O-PS-conjugated $MBP^{4xDQNAT}$ prepared in living *E. coli* cells using PCGT was used as a positive control. Each group was composed of six mice except for the PBS control group, which was composed of five mice. Mice were boosted on days 21 and 42 with identical doses of antigen. *Ft*LPS-specific IgG titers were measured by ELISA in serum collected on day -1, 35, 49, 63, and 70 following initial immunization. iVAX-derived bioconjugates elicited significantly higher levels of *Ft*LPS-specific IgG compared to compared to the PBS control group in serum collected on day 35, 49, and 70 of the study (**$p < 0.01$, Tukey-Kramer HSD). Values represent means and error bars represent standard errors of *Ft*LPS-specific IgGs detected by ELISA.

**Table S1. Cost analysis for iVAX reactions, Related to STAR Methods.** The total cost to assemble iVAX reactions is calculated below. A 1 mL iVAX reaction produces two 10 μg vaccine doses and can be assembled for $11.75. In the table, amino acid cost accounts for 2 mM each of the 20 canonical amino acids purchased individually from Sigma. Lysate cost is based on a single employee making 50 mL lysate from a 10 L fermentation, assuming 30 lysate batches per year and a 5-year equipment lifetime. Component source is also included in the table if it is available to purchase directly from a supplier. Homemade components cannot be purchased directly and must be prepared according to procedures described in the Methods section.

| Component | Cost ($/mL rxn) | Supplier | Product No |
|---|---|---|---|
| $Mg(Glu)_2$ | <0.00 | Sigma | 49605 |
| $NH_4Glu$ | <0.00 | MP | 02180595 |
| KGlu | <0.00 | Sigma | G1501 |
| ATP | 0.01 | Sigma | A2383 |
| GTP | 0.27 | Sigma | G8877 |
| UTP | 0.23 | Sigma | U6625 |
| CTP | 0.20 | Sigma | C1506 |
| Folinic acid | 0.02 | Sigma | 47612 |
| tRNA | 0.21 | Roche | 10109541001 |
| Amino acids | <0.00 | homemade | |
| PEP | 1.79 | Roche | 10108294001 |
| NAD | 0.07 | Sigma | N8535-15VL |
| CoA | 0.34 | Sigma | C3144 |
| Oxalic acid | <0.00 | Sigma | P0963 |
| Putrescine | <0.00 | Sigma | P5780 |
| Spermidine | <0.00 | Sigma | S2626 |
| HEPES | <0.00 | Sigma | H3375 |
| $MnCl_2$ | <0.00 | Sigma | 63535 |
| DDM | 0.36 | Anatrace | D310S |
| Plasmid | 0.88 | homemade | |
| Lysate | 7.37 | homemade | |
| **Total** | **11.75** | **$/mL rxn** | |
| | **5.88** | **$/dose** | |

**Table S2. Primers used to generate CLM24 Δ*lpxM*, Related to STAR Methods.** Primers used to construct and verify the CLM24 Δ*lpxM* strain are listed below. KO primers were used for amplification of the kanamycin resistance cassette from pKD4 with homology to *lpxM*. Seq primers were used for colony PCRs and sequencing confirmation of knockout strains.

| Primer Name | DNA Sequence (5' to 3') |
|---|---|
| *lpxM* KO for | TACACTATCACCAGATTGATTTTTGCCTTATCCGAAACTGGAAAAGCATGGTGTAGGCTGGAGCTGCTTC |
| *lpxM* KO rev | GCGAAGGCCTCTCCTCGCGAGAGGCTTTTTTATTTGATGGGATAAAGATCCATATGAATATCCTCCTTAGTTCCTATTC |
| *lpxM* seq for | AGTACCGGCTTTTTTTATTTGG |
| *lpxM* seq rev | CTAATACCACGCGTATTTTAACG |

**Table S3. Plasmids used in this study, Related to STAR Methods.**

| Plasmid | Description | Source |
|---|---|---|
| pSF-*Cj*PglB | *C. jejuni* PglB with a C-terminal 1xFLAG epitope tag in pSF, a modified pBAD expression vector | (Ollis et al., 2014) |
| pGAB2 | *F. tularensis* O-PS antigen gene cluster in pLAFR1 | (Cuccui et al., 2013) |
| pMW07-O78 | *E. coli* O78 antigen gene cluster in pMW07 | (Celik et al., 2015) |
| pJHCV32 | *E. coli* O7 antigen gene cluster in pVK102 | (Valvano and Crosa, 1989) |
| pKD46 | Encodes λ red system for recombineering | (Datsenko and Wanner, 2000) |
| pKD4 | Encodes kanamycin resistance cassette with upstream and downstream FRT sites | (Datsenko and Wanner, 2000) |
| pCP20 | Encodes *flp* for Flp-FRT recombination | (Datsenko and Wanner, 2000) |
| pSF-*Cj*PglB-LpxE | *C. jejuni* PglB with a C-terminal 1xFLAG epitope tag and *F. tularensis* LpxE in pSF | This work; Addgene 128389 |
| pJL1-sfGFP$^{217\text{-}DQNAT}$ | Superfolder green fluorescent protein variant modified after residue T216 with 21 amino acid insertion containing the *C. jejuni* AcrA N123 glycosylation site but with an optimal DQNAT glycosylation sequence and a C-terminal 6xHis tag | (Jaroentomeechai et al., 2018) |
| pJL1-sfGFP$^{217\text{-}AQNAT}$ | Same as pJL1 sfGFP$^{217\text{-}DQNAT}$, but with an AQNAT glycosylation sequence that is not modified by *Cj*PglB | (Jaroentomeechai et al., 2018) |
| pJL1-MBP$^{4xDQNAT}$ | *E. coli* maltose binding protein with a C-terminal 4xDQNAT glycosylation tag and a 6xHis tag in pJL1, a T7-driven *in vitro* expression vector | This work; Addgene 128390 |
| pJL1-PD$^{4xDQNAT}$ | *H. influenzae* protein D with a C-terminal 4xDQNAT glycosylation tag and a 6xHis tag in pJL1 | This work; Addgene 128391 |
| pJL1-PorA$^{4xDQNAT}$ | *N. meningitidis* PorA porin protein with a C-terminal 4xDQNAT glycosylation tag and a 6xHis tag in pJL1 | This work; Addgene 128392 |
| pJL1-TTc$^{4xDQNAT}$ | Fragment C domain of *C. tetani* toxin with a C-terminal 4xDQNAT glycosylation tag and a 6xHis tag in pJL1 | This work; Addgene 128393 |
| pJL1-TTlight$^{4xDQNAT}$ | Light chain variant of *C. tetani* toxin containing an inactivating E234A mutation in the enzyme active site with a C-terminal 4xDQNAT glycosylation tag and a 6xHis tag in pJL1 | This work; Addgene 128394 |
| pJL1-CRM197$^{4xDQNAT}$ | *C. diphtheriae* toxin variant with an inactivating G52E mutation in the enzyme active site with a C-terminal 4xDQNAT glycosylation tag and a 6xHis tag in pJL1 | This work; Addgene 128395 |
| pJL1-TT$^{4xDQNAT}$ | *C. tetani* toxin variant containing an inactivating E234A mutation in the enzyme active site with a C-terminal 4xDQNAT glycosylation tag and a 6xHis tag in pJL1 | This work; Addgene 128396 |
| pJL1-EPA$^{DNNNS\text{-}DQNRT}$ | *P. aeruginosa* exotoxin A containing a DNNNS glycosylation site at residue 242 and a DQNRT glycosylation site at residue 384 and a C-terminal 6xHis tag in pJL1 | This work; Addgene 128397 |

**Table S3 (continued). Plasmids used in this study, Related to STAR Methods.**

| Plasmid | Description | Source |
|---|---|---|
| pTrc99s-ssDsbA-MBP$^{4xDQNAT}$ | *E. coli* maltose binding protein with an N-terminal DsbA signal sequence for periplasmic translocation and a C-terminal 4xDQNAT glycosylation tag and a 6xHis tag in pTrc99s | This work; Addgene 128398 |
| pTrc99s-ssDsbA-PD$^{4xDQNAT}$ | *H. influenzae* protein D with an N-terminal DsbA signal sequence for periplasmic translocation and a C-terminal 4xDQNAT glycosylation tag and a 6xHis tag in Trc99s | This work; Addgene 128399 |
| pTrc99s-ssDsbA-PorA$^{4xDQNAT}$ | *N. meningitidis* PorA porin protein with an N-terminal DsbA signal sequence for periplasmic translocation and a C-terminal 4xDQNAT glycosylation tag and a 6xHis tag in pTrc99s | This work; Addgene 128400 |
| pTrc99s-ssDsbA-TTc$^{4xDQNAT}$ | Fragment C domain of *C. tetani* toxin with an N-terminal DsbA signal sequence for periplasmic translocation and a C-terminal 4xDQNAT glycosylation tag and a 6xHis tag in pTrc99s | This work; Addgene 128401 |
| pTrc99s-ssDsbA-TTlight$^{4xDQNAT}$ | Light chain variant of *C. tetani* toxin containing an inactivating E234A mutation in the enzyme active site with an N-terminal DsbA signal sequence for periplasmic translocation and a C-terminal 4xDQNAT glycosylation tag and a 6xHis tag in pTrc99s | This work; Addgene 128402 |
| pTrc99s-ssDsbA-CRM197$^{4xDQNAT}$ | *C. diphtheriae* toxin variant with an inactivating G52E mutation in the enzyme active site with an N-terminal DsbA signal sequence for periplasmic translocation and a C-terminal 4xDQNAT glycosylation tag and a 6xHis tag in pTrc99s | This work; Addgene 128403 |
| pTrc99s-ssDsbA-EPA$^{DNNNS-DQNRT}$ | *P. aeruginosa* exotoxin A containing a DNNNS glycosylation site at residue 242 and a DQNRT glycosylation site at residue 384 with an N-terminal DsbA signal sequence for periplasmic translocation and a C-terminal 6xHis tag in pTrc99s | This work; Addgene 128404 |

**Table S4. Antibodies and antisera used in this study, Related to STAR Methods.**

| Target | Source | Dilution |
|---|---|---|
| Rabbit pAb to 6xHis epitope tag | Abcam | 1:7500 |
| Mouse mAb FB11 to *F. tularensis* LPS | Abcam | 1:5000 |
| Rabbit pAb to *E. coli* O78 antigen | Abcam | 1:2500 |
| Rabbit pAb to *C. diphtheriae* toxin | Abcam | 1:2000 |
| Rabbit pAb to *C. tetani* toxin | Abcam | 1:2000 |
| Goat anti-rabbit IgG IR dye 680 | LI-COR | 1:15000-1:10000 |
| Goat anti-rabbit IgG IR dye 800 | LI-COR | 1:15000-1:10000 |
| Goat anti-mouse IgG IR dye 800 | LI-COR | 1:15000-1:10000 |
| Goat anti-mouse IgG HRP | Abcam | 1:25,000 |
| Goat anti-mouse IgG1 HRP | Abcam | 1:25,000 |
| Goat anti-mouse IgG2a HRP | Abcam | 1:25,000 |
